# Stochastic Phylogenetic Models of Shape

**DOI:** 10.1101/2025.04.03.646616

**Authors:** Sofia Stroustrup, Morten Akhøj Pedersen, Frank van der Meulen, Stefan Sommer, Rasmus Nielsen

## Abstract

Phylogenetic modeling of morphological shape in two or three dimensions is one of the most challenging statistical problems in evolutionary biology. As shape data are inherently correlated and non-linear, most naïve methods for phylogenetic analysis of morphological shape fail to capture the biological realities of evolving shapes. In this study we propose a novel framework for evolutionary analysis of morphological shape which facilitates stochastic character mapping on landmark shapes. Our framework is based on recent advances in mathematical shape analysis and models the evolution of shape as a diffusion process that accounts for the evolutionary correlation between nearby landmarks. The diffusion process we consider is parametrized in terms of meaningful parameters describing the evolutionary rate and the degree of spatial autocorrelation among landmarks. The framework we propose assumes that the phylogenetic tree is fixed and uses a Metropolis-Hastings Markov Chain Monte Carlo sampling scheme for inferring ancestral shapes and parameters of the model. We evaluate the new inference algorithm using simulations and show that the method leads to improved estimates of the shape at the root and well-calibrated credible sets of shapes at internal nodes. In addition, we also compare the diffusion parameter describing the degree of spatial autocorrelation to an existing metric of integration and find that they quantify integration in a shape in a similar way. To illustrate the method, we also apply it to a previously published data set of butterfly wing images.

## Introduction

A core objective of phylogenetics is to understand and analyze the evolution of measurable traits along the edges of a phylogeny. One common application is character mapping in which the ancestral states of a trait are mapped to internal nodes in a phylogeny, when only the states at the leaf nodes have been observed (Nielsen, 2002; Huelsenbeck et al., 2003; Bollback, 2006; Goolsby, 2017). Mapping of traits on a phylogeny forms the foundation of many downstream statistical analyses, for example the detection of evolutionary correlations among traits or identification of correlations between environmental and morphological variables (Felsenstein, 1985). Traditionally, character mapping is done using parsimony for discrete variables, but this approach has the disadvantage of being biased towards inferring as little change as possible and not representing statistical uncertainty (Nielsen, 2002). This has led to a focus on stochastic character mapping that circumvents these challenges by using stochastic models of character evolution to represent uncertainty (Nielsen, 2002; Huelsenbeck et al., 2003). Such approaches can also naturally be applied to continuous traits using stochastic processes on a continuous state space, including Brownian motion models (Martin and Weber, 2024).

Modern phylogenetics relies on a suite of stochastic models of character evolution that extend the simple Brownian motion model, including the Ornstein-Uhlenbeck process (Hansen, 1997; Butler and King, 2004) and the Early Burst model (Blomberg et al., 2003; Harmon et al., 2010). These methods are applicable to traits represented by a few variables, such as weight, height, or blood pressure (Bartoszek et al., 2012; Mitov et al., 2020). However, it is more challenging to apply these methods when there are multiple correlated traits. Reliable inference of the evolutionary covariance matrix, with *p*^2^ elements for *p* traits, requires data from many species (*n* ≫ *p*; where *n* is the number of species) and sufficient evolution along the edges to allow identifiability. Therefore, a common practice has been to reduce the dimensionality of the data using principal component analysis (PCA) and then use the major principal components as independently evolving variables in down-stream analysis (see for example Monteiro and Nogueira (2010); Hunt (2013) and general recommendations for this type of analysis are described in Monteiro (2013)). However, this approach has some unsolved disadvantages, including possible biases introduced by the linear transformation performed by the PCA (Revell, 2009; Uyeda et al., 2015) and the lack of biological interpretability when assuming principal components are the units of evolution rather than the original characters. A number of other solutions to the large *p* small *n* problem has been proposed including the use of distance-based methods for phylogenetic regression (Adams, 2014a,b), pairwise composite likelihood for estimation of parameters (Goolsby, 2016), and penalization (Clavel et al., 2019).

Morphological shape is a particularly challenging problem. The shape itself is infinite dimensional, and even if represented by landmarks it will typically be very high-dimensional with each trait (landmark) being highly correlated with other traits (adjacent landmarks). In geometric morphometrics, a shape has been defined following the work of Kendall as ‘all the geometrical information that remains when location, scale and rotational effects are removed from an object’ (Dryden and Mardia, 2016). In practice, this notion of a shape is reflected when Procrustes alignment (Gower, 1975) is used to pre-process the data as Procrustes alignment superimposes the data by means of rotation, translation, and scaling. The current state-of-the-art in phylogenetic analyses of shape data involves 1) pre-processing the shape data using Procrustes alignment and 2) applying the same stochastic models as for non-shape data (such as Brownian motion, Early Burst or OrnsteinUhlenbeck) to the Procrustes aligned data, possibly after a phylogenetic principal component analyses (PCA) that attempts to reduce landmarks to a set of approximately independently evolving variables. Applying the stochastic models typically used in stochastic character mapping (e.g. Brownian motion and Ornstein-Uhlenbeck) to model the evolution of morphological shape has some unsolved disadvantages, namely that the state space of these models will include configurations of landmarks that do not look like morphological shapes in the sense that the outline of a shape (denoted by landmarks) might cross itself and the landmarks overlap. We illustrate this disadvantage by considering what happens when we simulate data under these models. If we let each landmark evolve as independent Brownian motions the collection of points will likely evolve into something that cannot represent biological shapes, as illustrated in Figure 2a. One way to handle this is to fit an empirical covariance matrix to a data set of landmark shapes and model the shape evolution by a Brownian motion with this covariance, thus enforcing the observed empirical covariance between landmarks (as described in Polly (2004)). From a shape perspective, this approach is valid only for evolutions over ‘small’ periods of time relative to the distance between landmarks, as also mentioned by Polly (2004). The reason for this is that such a Brownian motion model with a fixed covariance allows for landmarks to cross each other, as illustrated in Figure 2b. Although the approach described in Polly (2004) lead to biologically reasonable shapes under short timescales it still does not solve the underlying problem of modeling shape. This emphasizes the need for new stochastic models of shape and raises the question whether it is possible to use stochastic shape models for phylogenetic analysis.

D’Arcy Thompson argued in the early 20th century that morphological evolution can be described as a deformation of the coordinate system (Thompson, 1917). In this paper we will explore the development of stochastic models of shape in the spirit of D’Arcy Thompson that arise from recent advances in stochastic shape analysis (Arnaudon et al., 2019). These models have the property that the stochastic process at all time points looks like a shape if the initial state is a shape, i.e. the state space of the process is the set of all supported shapes. Furthermore, when using landmarks, they take advantage of the distance between landmarks to model the correlation structure parametrically. This solves the *p > n* problem. Increasing the number of landmarks does not lead to identifiability issues as the process is parameterized in terms of deformations in a 2 or 3-dimensional coordinate systems. In other words, as the number of landmarks increases, the number of parameters stay the same, thereby we avoid the *p > n* problem which is normally encountered when modelling shape data. Technically, this is done by letting the covariance of each step of the process depend on the distances between the landmarks at any given time. We can think of the process as a shape-aware Brownian motion. In line with the general recommendations in the field of geometric morphometrics (see discussion in Adams and Collyer (2017)) the stochastic model we apply is rotation invariant as the distance between landmarks does not change when rotating the data. This means that parameter and ancestral state estimates do not depend on the orientation of the data.

We illustrate the use of the shape process for stochastic character mapping, i.e. for the statistical inference of ancestral shapes on the phylogeny. In this manuscript we assume the phylogeny to be given and fixed. Standard algorithms for stochastic character mapping assume that transition probabilities or transition densities of the stochastic process can be calculated analytically or efficiently numerically. Unfortunately, that is not true for the shape process we propose. We will instead use the approach introduced by van der Meulen et al. (2025) where paths are simulated under an approximative process that provides weighted samples of the conditioned process. We then use Markov Chain Monte Carlo (MCMC) methods to sample from the joint posterior of the shape process on the phylogeny and the unknown parameters. This provides a complete framework for Bayesian inference of ancestral shapes and parameters governing the evolutionary shape process. This approach provides credible sets of ancestral shapes that incorporate uncertainty in the parameters of the shape process. We evaluate the method using simulations and also show an application to real data of butterfly wing images from Chazot et al. (2021).

In Figure 1 we provide an overview of the proposed method for shape-aware stochastic character mapping. In order to use our method we need a phylogeny with shapes associated to all leaf nodes (Fig. 1a). The shapes should be defined in terms of landmarks as shown in Figure 1b, although the inference framework in theory can be generalized to full shapes (infinite-dimensional processes). We then assume a diffusion model that models the evolution of shape (Fig. 1c and 1d). The diffusion model is parametrized in terms of meaningful parameters describing the evolutionary rate or the degree of autocorrelation among landmarks. The stochastic shape model is fully specified in the section *“Model and Parameters”*. We adopt a Bayesian inference procedure, which allows us to infer the joint posterior distribution over the evolutionary rate, the degree of autocorrelation among landmarks and the ancestral states. This is illustrated in Figure 1e and Figure 1f.

**Figure 1.**
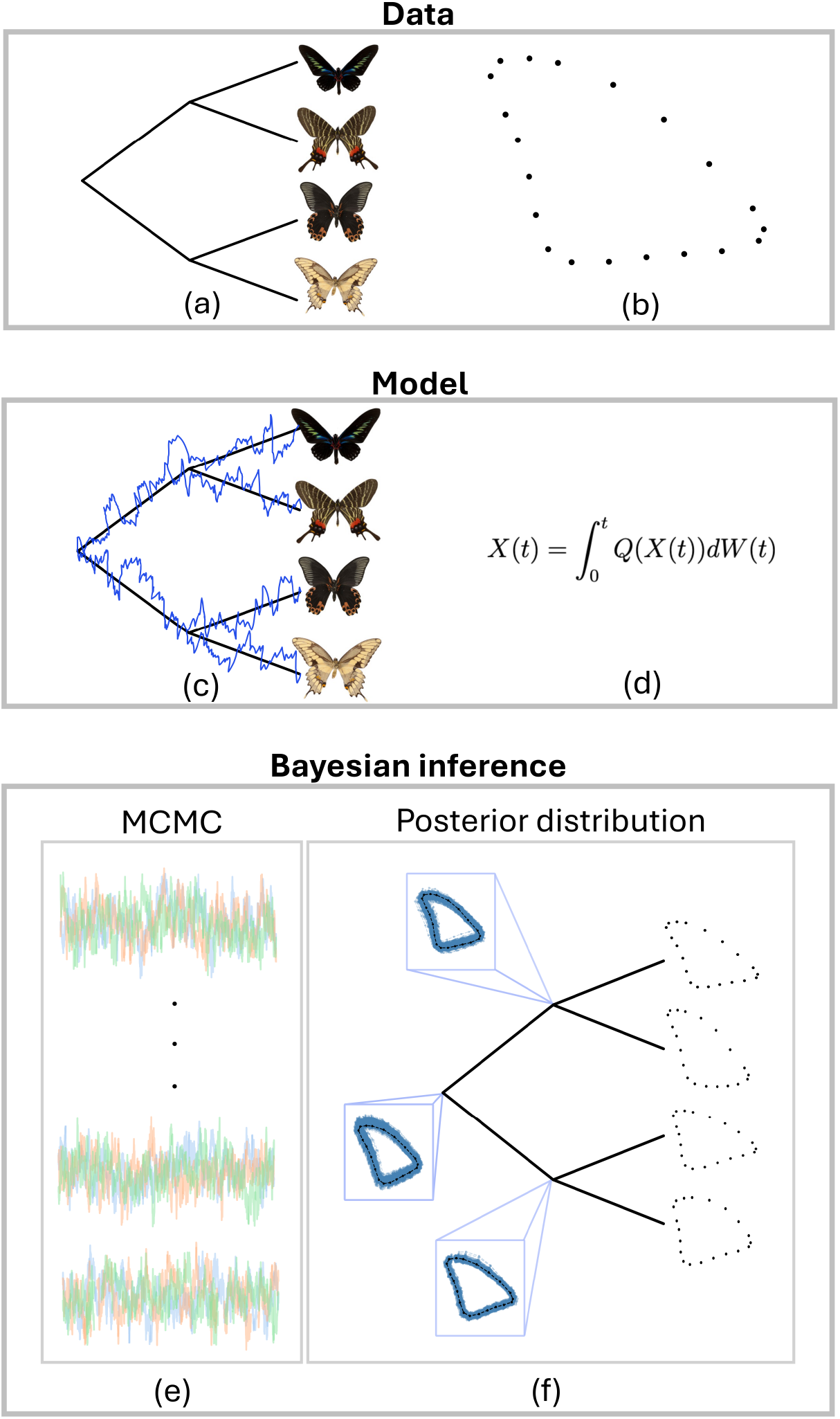
An overview of the proposed method. (a) An example of a data set. We assume that the phylogenetic tree and the shapes at the tips are given. (b) A butterfly forewing outlined using landmarks. (c) The stochastic shape process evolving on the phylogeny. (d) The infinite-dimensional formulation of the stochastic shape process. (e) Trace plots obtained from the MCMC. (f) Posterior distribution over ancestral states. Images of butterflies were obtained from GBIF and distributed under CC0 1.0 or CC BY-NC 4.0 (see Supplementary Material Fig. S1).

## Materials and Methods

In this section we describe the shape model and the inference methodology used to infer the posterior distribution. For completeness, we describe the shape model and inference strategy at different levels of mathematical detail. We refer to subsection *“Model and Parameters”* and subsection *“General Ideas on the Methodology”* for a general introduction to the model and the inference methodology used. Finally, we also describe empirical data sets, simulation strategies and existing methods used in the experiments presented in section *“Results”*.

### Model and Parameters

Following Sommer et al. (2021), we model shape dynamics as a stochastic process satisfying the following three principles.

A. The covariance between landmarks of a shape *x* depends only on the distances between the landmarks; nearby landmarks will have higher correlation compared to landmarks that are further apart.
B. We model the shape evolution as a diffusion process whose diffusion matrix is a square root of the covariance matrix. The diffusion matrix is thus state dependent – it depends on the distances between landmarks at the current time point.
C. The stochastic evolution of the shape is *diffeomorphic* (explained below).

We briefly mention how these principles differ from the stochastic processes which are currently being used to model the evolution of shapes such as the Brownian motion model, the Early Burst model, and the Ornstein-Uhlenbeck process. Point A): our covariance follows a parametric form which depends on the distances between landmarks whereas the standard models have a non-parametric covariance. Point B): The diffusion matrix changes as the shape changes, whereas it is fixed in the Brownian motion model and OrnsteinUhlenbeck process and time dependent in the Early Burst model. Point C): A diffeomorphism *φ* on ℝ^*d*^ is a map from ℝ^*d*^ to ℝ^*d*^ which is differentiable and whose inverse map exists and is differentiable. That the shape evolution is diffeomorphic i.e. modelled using invertible and differentiable maps, ensures that, for arbitrarily long time periods, landmarks will not cross. In fact, landmarks distributed along non-self-intersecting curves or surfaces will preserve this property which is arguably crucial when modeling shape evolution. This is the case as the map is one-to-one, i.e. invertible, at each time point which ensures that two distinct points cannot be mapped to the same point. If two landmarks were to cross, they would be overlapping at the crossing time, and this is not possible when we use diffeomorphisms to model shape evolution. A closely related property is, furthermore, that the diffeomorphic dynamics are independent of the choice of landmarks, in a sense explained below. This is in contrast to the Brownian motion model which is valid only for a prespecified number of landmarks. Diffeomorphisms provide a precise mathematical formulation of what D’Arcy Thompson described as ‘deformations’ in his pioneering work (Thompson, 1917) inspiring geometric morphometrics.

We consider our stochastic model to be an analogue of a standard Brownian motion, an evolutionary null model, for shapes. It contains no drift, and the diffusion term is of a simple form that ensures the crucial diffeomorphic property, without imposing further assumptions.

In the following section we define the process. It can be described at different levels of mathematical abstraction, in terms of either finite or infinite dimensional objects. The finite-dimensional formulation in section *“Finite-dimensional Formulation of the Shape SDE”* is mathematically most accessible. This formulation is also the one first introduced in Arnaudon et al. (2019). Our numerical results are obtained by a version of this finite-dimensional formulation. The infinite-dimensional formulation is described in Appendix 1. This requires more mathematical machinery, which is outside the scope of this paper, but it shows that the finite-dimensional formulation is a discretized version of a mathematical object that does not depend on the choice of discretization. Further mathematical details on stochastic shape processes can be found in Sommer et al. (2025).

#### Finite-dimensional Formulation of the Shape SDE

We represent a shape as a set of *k* points (landmarks) *q*_1_, …, *q*_*k*_ in ℝ^*d*^, for *d* = 2 or 3. Equivalently, such a shape can be considered a single (*d* · *k*)-dimensional vector, *x* ∈ ℝ^*d*·*k*^. Let *p*_1_, …, *p*_*J*_ be *J* points on a grid in *D* ⊂ ℝ^*d*^, illustrated as dots on Figures 2c and 2d. To each grid point *p*_*j*_, *j* = 1 … *J*, we associate a kernel function 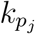: *D* → ℝ centered at *p*_*j*_. In this work, we choose a scaled Gaussian density

**Figure 2.**
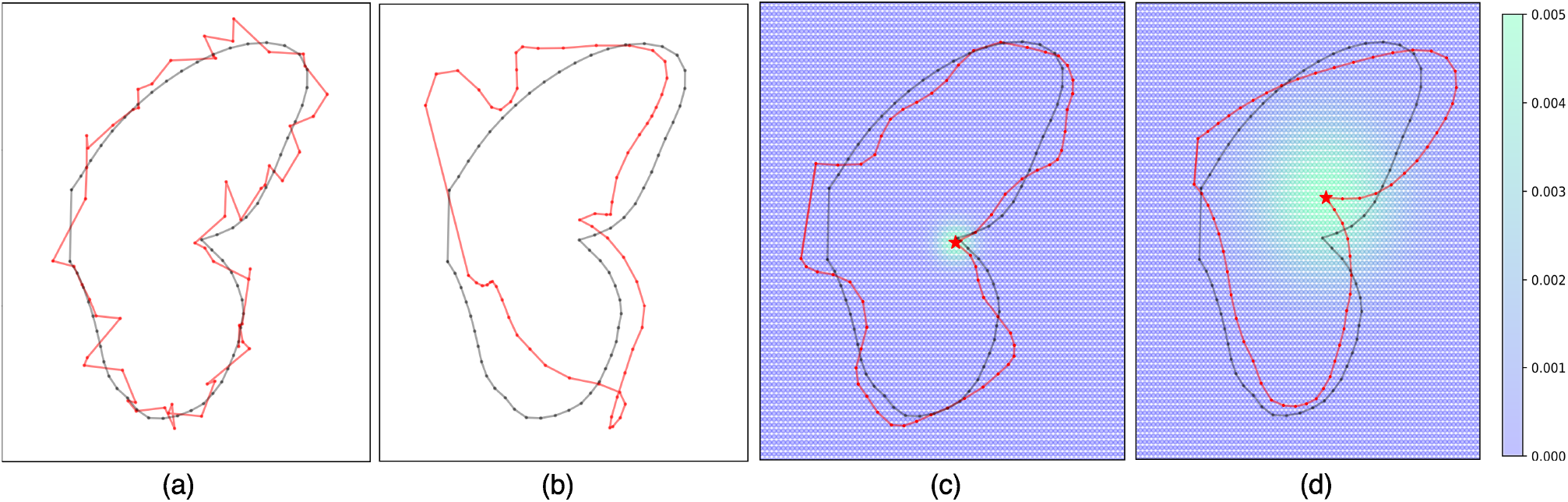
In each figure, the black dots show initial positions of 59 landmarks. The red dots show landmark positions at time *t* = 1 for each of the following 4 stochastic processes. (a) Independent Brownian motions. (b) A 118-dimensional Brownian motion with (time-independent) covariance matrix (as defined in (4)) evaluated at the initial wing configuration, using kernel parameters *σ* = 0.1, *α* = 0.05. (c) A solution to the SDE given in (3) with kernel parameters *σ* = 0.1 and *α* = 0.005. Dots in the background shows placement of grid points *p*_*j*_, *j* = 1 … 6400, coloured by the values of the kernel 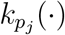 evaluated at the starshaped landmark. (d) Identical to c except for a higher value of *σ* = 0.4.

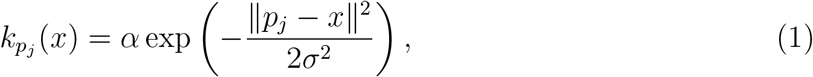

where *α* is the *amplitude* and *σ* the standard deviation, or *width*, of the kernel. The value 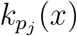 is largest for *x* = *p*_*j*_ and decreases with the distance between *x* and *p*_*i*_. To each grid point *p*_*j*_ we associate an ℝ^*d*^-valued standard Brownian motion *W* ^*j*^. We assume the landmark at initial location *q*_1_ evolves according to the stochastic differential equation (SDE):

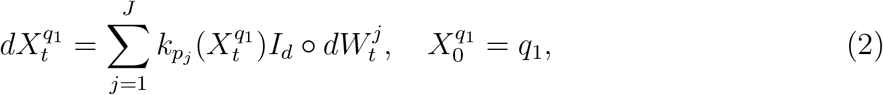

where *I*_*d*_ denotes the *d × d* identity matrix and ‘∘’ denotes a Stratonovich stochastic integral, which can equivalently be converted to an Itô integral by adding a drift correction term (see, e.g. Oksendal (2013)). The SDE is in fact an integral equation to be interpreted as

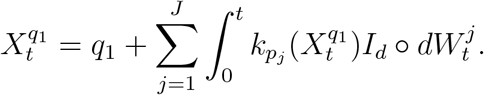

The difference between Itô and Stratonovich is caused by either evaluating the integrand at the midpoint or left-point in Riemann-sum approximations. At each time *t*, the solution 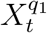 moves in the direction of the sum of all Brownian motions 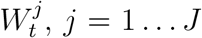, whose kernel value 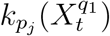 is non-negligible. As detailed in Kunita (1990), the SDE induces a *stochastic flow of diffeomorphisms*, meaning that at every time *t* the map from the initial point *q*_1_ to the state at time *t*, i.e. 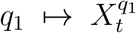, is a diffeomorphism. The diffeomorphic property ensures the following: if we let *q*_1_, …, *q*_*k*_ be initial landmarks along a closed, non-self-intersecting curve or surface in *D* ⊂ ℝ^*d*^, then 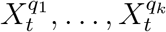, will be points on a stochastically deformed curve or surface that is still closed and non-self-intersecting. Furthermore, this stochastically deformed curve can be represented by an arbitrary number of landmarks, without changing the underlying dynamics, as detailed in the next paragraph.

The simultaneous dynamics of multiple points, i.e. landmarks, under the SDE (2) is given by the system of SDE’s, all driven by the same noise fields *W* ^*j*^, *j* = 1 … *J*:

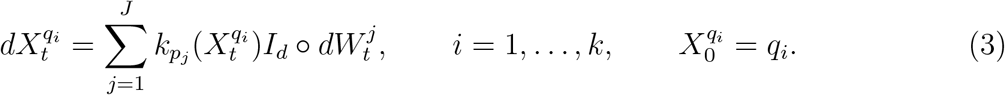

This system can equivalently be formulated as a single SDE on ℝ^*d*·*k*^,

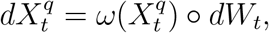

where *W* is a (*d* · *J*)-dimensional standard Brownian motion, *q* = (*q*_1_, …, *q*_*k*_) ∈ ℝ^*d*·*k*^ is the concatenated vector of landmarks of initial shape, and the diffusion matrix is given by the following block matrix

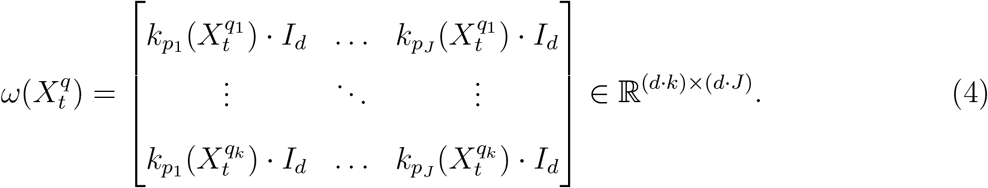

Note that the *k d*-dimensional equations in the SDE (3) are independent of each other, given a realization of the Brownian motions. However, the trajectories are correlated, since they are driven by the same Brownian motions. An important consequence of this is that, once the Brownian motions 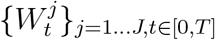 have been sampled up to time T, the dynamics of any arbitrary point in *D* is determined and can be computed.

The covariance of a shape at time *t*, i.e. the covariance between its landmarks, is computed from the diffusion matrix by 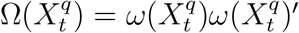. The covariance between coordinates of two different landmarks *i* and *m* is thus given by 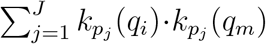, and the covariance is isotropic so that it is the same in all dimensions and along any direction. This means that landmarks which are far away from each other, relative to the kernel width, have small covariance. This point is illustrated further in the next paragraph. Note that the preceding discussion is the precise formulation of points A) and B) mentioned in section *“Model and Parameters”*.

In the remainder of this section, we elaborate on why solutions to this SDE yield the desired correlated motion of landmarks. Firstly, note that the amplitude *α* is a scaling factor of the kernel (1) and thus the Brownian motions, and thereby it affects only the speed of the trajectory. The parameter *α* can therefore be interpreted, in an evolutionary context, as a *rate* parameter. We explain the width parameter, *σ*, by considering the following SDE governing the difference between two landmark trajectories;

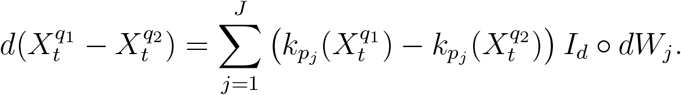

This SDE implies that if 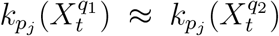 for each grid point *j* then the difference will be close to constant – i.e. the landmarks will move in parallel. 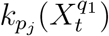 is close to 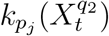 for all *j* if the landmarks 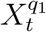 and 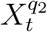 are close to each other *relative to the width* of the kernel. The more narrow the kernel is, the closer landmarks need to be in order for their displacements to be parallel, and vice versa. A wide kernel implies that landmarks further apart will be displaced parallelly. Thus, in geometric morphometrics terminology, *σ* roughly corresponds to a measure of overall ‘integration’ of the shape. See Figure 2c and 2d for an illustration of the effect of different kernel widths. Each figure shows the position of 59 landmarks at time 1 (red dots) after following the SDE (3) initialized in the configuration of a butterfly wing (black dots). The kernel width (standard deviation) is *σ* = 0.1 for Figure 2c and *σ* = 0.4 for Figure 2d. The amplitude is *α* = 0.005 in both cases. As can be seen from Figure 2, the shape that has evolved according to the SDE with the smallest kernel width (Fig. 2c) is more wiggly than the shape that has evolved according to the SDE with the largest kernel width (Fig. 2d). This is what we would expect as a larger *σ* parameter results in a larger kernel width, making the individual landmarks correlated over longer distances.

### Inference

#### General Ideas on the Methodology

In this paper we adopt the Bayesian point of view towards statistics. This means that we specify information about unknown parameters in the statistical model by probability distributions. In the proposed shape model, the unknown parameters are *α* and *σ*, the amplitude and standard-deviation of the kernel. In a second stage, we will additionally define a prior on the root shape, rather than assuming it to be fixed and known. For now, assume only *θ* := (*α, σ*) is unknown and we have specified a prior distribution on *θ*. The data we observe are the shapes at the leaf nodes, which we denote by 𝒟. If the likelihood of 𝒟, evaluated at *θ*, would be known in closed form the problem would be simple: denoting the likelihood by *L*(*θ*; 𝒟) and the prior density on *θ* by *π*, by Bayes’ theorem the posterior density at *θ* is given by *π*(*θ* | 𝒟) = *L*(*θ*; 𝒟)*π*(*θ*)*/* ∫ *L*(*θ*; 𝒟)*π*(*θ*)*dθ*. The integral over ℝ^2^ can easily be numerically approximated. As we detail below, *L*(*θ*; 𝒟) is intractable (it is not known in closed form), which poses a key difficulty that we need to work around.

There exists a recursive backward scheme to compute the likelihood, starting from the leaves back to the root vertex (a postorder traversal of the tree), see van der Meulen et al. (2025) and references therein. Within this scheme, which is a general version of Felsenstein’s algorithm, one computes partial (or fractional) likelihoods, i.e. likelihoods of subtrees, denoted by *h* in the following. In Felsenstein’s algorithm, the likelihood can be obtained directly. Similarly, latent data at internal nodes can be simulated conditionally on the states at the leaf nodes by first using Felsenstein’s algorithm to calculate partial likelihoods for all nodes. Then simulations can proceed from the root towards the leaves (pre-order traversal) using the simulated state at the parent node, and the partial likelihood at the child node, at each step. This can be incorporated into MCMC schemes for joint inference of the path of the process and parameters of the process (see for instance Nielsen (2001); Irvahn and Minin (2014)). While this scheme has been applied to discrete characters, and recently also under simple Brownian motion process (Martin and Weber, 2024), a similar algorithm can be used for general diffusion processes (van der Meulen et al., 2025). The computed partial likelihoods can be used to sample paths from the shape process, conditional on the shapes at the leaf vertices. In fact, it turns out that the conditioned process has the same diffusivity as the (unconditional) forward shape process, but with an additional drift superimposed that depends on the computed partial likelihoods. Intuitively, the partial likelihoods induce a reweighting of each path on the tree, where paths that match the data at the leaf vertices get higher probability and paths that contradict the data get probability zero.

The key difficulty is that the modified Felsenstein algorithm requires solving the Kolmogorov backward equation on each edge. This is a partial differential equation which typically cannot be solved in closed form. Backward Filtering Forward Guiding (BFFG) resolves this problem using two key ideas:

1. The backward recursion is computed under a simplification of the forward model such that its computations become tractable. In other words, we ensure we can easily solve the Kolmogorov backward equation on each edge. The partial likelihoods obtained under this simplification are denoted by *g* and referred to as guiding functions.
2. We can use the guiding functions just like the partial likelihoods to reweight paths. The resulting process is called the guided process. It is constructed to resemble the true conditioned process. The discrepancy between the guided process and true conditioned process can be quantified using the likelihood ratio between the probability measures of both processes. Next, the Metropolis-Hastings algorithm can be invoked to sample from the joint distribution of the parameters and the latent shape process on the tree, conditional on the data 𝒟.

#### Mathematical Preliminaries

The probabilistic dynamics of the shape process can be described in three ways:

A. The Stratonovich SDE that we present. This approach is amenable to both interpretation and stochastic simulation.
B. Transition probabilities, which specify the distribution of *X*_*t*_, conditional on *X*_*s*_ = *x* when *s < t*.
C. The infinitesimal generator (IG) of the process. For the *d*-dimensional Itô-diffusion process *dX*_*t*_ = *b*(*X*_*t*_)*dt* + *ω*(*X*_*t*_)*dW*_*t*_ the infinitesimal generator is defined as the differential operator that acts on functions *f* : ℝ^*d*^ → ℝ as follows

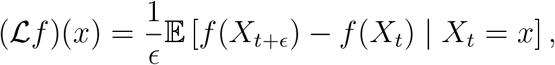

defined for those *f* for which the limit exists (these functions *f* are then said to be in the domain of the IG). In that case, one can express this operator in terms of the drift and diffusivity of the SDE:

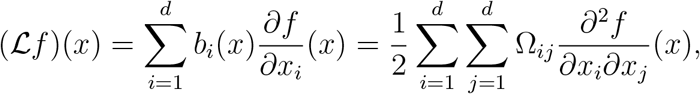

where Ω = *ωω*′. This operator captures how the process locally behaves.

It turns out that if the domain of the IG is sufficiently large, one can switch between (A), (B), and (C). Suppose transition densities *p* exist, i.e. for *t < T* and *A* ⊆ ℝ^*d*^, ℙ (*X*_*T*_ ∈ *A* | *X*_*t*_ = *x*) = ∫_*A*_ *p*(*t, x*; *T, y*)*dy*. Fixing *T* and *y*, define *u*_*t*_(*x*) = *p*(*s, x*; *T, y*). It is a well-known result from probability theory of stochastic processes that *u* satisfies the Kolmogorov backward equation:

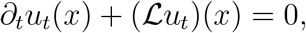

where we denote 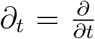. This equation can be written more compactly as (∂_*t*_ + ℒ)*u* = 0. This equation provides a link between descriptions (B) and (C). In the recursive scheme we use in our methodology, this equation will be solved backwards provided the “initial” condition *u*_*T*_.

#### General Setup

We consider the stochastic process *X* taking values in ℝ^*d*^ evolving on a directed tree. We assume the value at the root node, *x*_0_, is fixed and known, a condition we will relax later. When the process splits, it does so conditionally independently. The process is only observed at its leaf nodes. Denote the set of leaf nodes by 𝒱 and the values observed by *x*_*V*_ := {*x*_*v*_, *v* ∈ 𝒱}. On each edge connecting internal nodes, the process evolves according to the SDE specified in (3).

In the following, the set of internal nodes is defined to be the set of all nodes except for the root node and leaf nodes. For any node *s* other than the root node we denote by pa(*s*) its unique parent node and by ch(*s*) the set of its child nodes. For a subset of nodes *S* we denote *x*_*S*_ = {*x*_*s*_, *s* ∈ *S*}. An ‘internal edge’ is an edge where the shape diffusion process evolves. Such an edge, connecting nodes *s* and *t*, is denoted by *e* = (*s, t*). The full set of internal edges is denoted by ℰ. We assume that the diffusion process evolves on internal edge *e* for time *τ*_*e*_ according to the SDE specified in (3).

#### Likelihood and Conditioned Process

In this section we will show how the likelihood of *x*_𝒱_ can be computed recursively and characterise the process *X*, conditional on *x*_𝒱_. If the process would take values in a finite set and evolve in discrete steps between nodes connecting an edge or if the transition probability is known in closed form (which is the case in standard stochastic character mapping), Felsenstein’s algorithm provides a simple recursive way to compute the likelihood efficiently using a postorder traversal.

With small adaptations, Felsenstein’s algorithm applies to our setting as well, although we cannot compute the transition probability in closed form. See van der Meulen et al. (2025); van der Meulen (2022); Stoltz et al. (2021) for example. Let *u* be an internal node of the tree and define 𝒱(*u*) to be the set of leaf nodes in the subtree with root node *u*. Define the partial likelihood at node *u* by

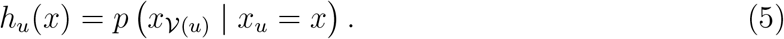

This is the density of the observations in the subtree, conditional on the process being in state *x* at node *u*. We assume that this density exists. Note that this definition implies that the likelihood is given by *h*_0_(*x*_0_), if conditioned on known root. A modified version of Felsenstein’s algorithm where the transition probabilities are obtained using the Kolmogorov backward equation is determined by the following rules.

1. Any leaf node *v* sends a *message m*_(pa(*v*),*v*)_ to its parent, defined by *m*_(pa(*v*),*v*)_(*x*) = *p*(*x*_*v*_ | *x*_pa(*v*)_ = *x*).
2. On an internal edge *e* = (*s, t*), node *t* sends to *s* the *message m*_*e*_ := *h*_*e*,0_, where *x* ↦ *h*_*e*,0_(*x*) is obtained from backward solving the Kolmogorov backward equation:

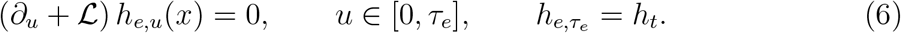

Here ℒ is the infinitesimal generator of *X*, introduced under (C) in section *“Mathematical Preliminaries”*. If fully written out, (6) is a partial differential equation. Its precise form is of minor importance at this stage. Note that there is a slight abuse of notation, where the subscript on *h* can refer to either the node or time instance on an internal edge.
3. When an internal node *s* has received messages from *all* of its children nodes, we set *h*_*s*_(*x*) = Π_*t*∈ch(*s*)_ *m*_*s,t*_(*x*). In this step, that we call *fusion*, all incoming messages at the parent node *s* are combined.

In other words, this is a more general version of Felsenstein’s algorithm than those that have hitherto been introduced (such as Mitov et al. (2020)) as it also works when transition densities cannot be computed in closed form. In practice this means that this general version of Felsenstein’s algorithm also works for non-Gaussian stochastic models.

Once *h* has been computed back to the root-node, the process *X* that is conditioned on its values at the leaf nodes –denote this by *X*^⋆^– satisfies the same SDE as the forward process, but with the additional drift term

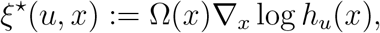

where Ω(*x*) = *ω*(*x*)*ω*(*x*)′. Thus, on the edge *e* = (*s, t*), *X*^⋆^ solves the SDE

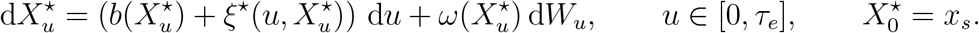

Here *b* is the Itô-Stratonovich correction term from converting the SDE in (3) to its Itôform. If it were possible to compute *h* in closed form in the modified Felsenstein algorithm (1)–(3), we would have an expression for the likelihood and the conditioned process could be simulated using a numerical SDE-solver for discretising *X*^⋆^.

#### Guiding

Unfortunately, (6) does not admit a closed form solution. So neither the likelihood nor the conditioned process dynamics can be computed. The crucial idea underlying the guided proposal framework considered in Schauer et al. (2017); van der Meulen et al. (2025); Mider et al. (2021), consists of finding substitutes to the messages *h*, which we denote by *g* such that the computations in steps (1)–(3) become tractable.

1. We assume at a leaf node *x*_*v*_ | *x*_pa(*v*)_ ~ *N* (*x*_pa(*v*)_, Γ) i.e. we assume Gaussian measurement noise. As the noise we assume is Gaussian the corresponding message *m*_*e*_, with *e* = (pa(*v*), *v*), can be written as log 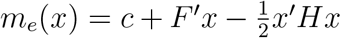, with *c* = log *φ*(*v*; 0, Γ), *F* = Γ^−1^*v* and *H* = Γ^−1^.
2. We choose simplified dynamics of the process *X*, say 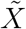 with infinitesimal generator 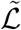, such that we can solve equation (6) for this process. The choice 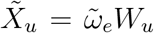 will ensure this. We detail the choice of 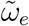 below. Let *g* solve

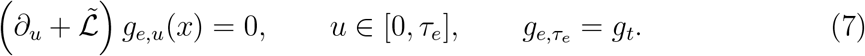

It is shown in Mider et al. (2021) that if *g*_*t*_ can be written as log 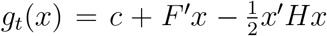, then for *all u* ∈ [0, *τ*_*e*_] the solution to (7) can be written as

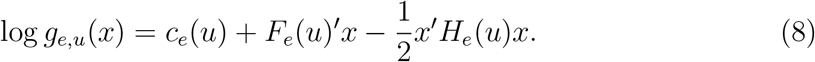

Therefore, we can represent the solution at any time *u* ∈ [0, *τ*_*e*_] by the triplet (*c*_*e*_(*u*), *F*_*e*_(*u*), *H*_*e*_(*u*)). Note that there is no direct relation between *h* and *H* and that we simply use the same notation as in Mider et al. (2021) to denote the triplet.
3. By the previous two steps, messages can always be parametrised by a triplet (*c, F, H*). Suppose on the edge *e*, the message *m*_*e*_ is parametrised by (*c*_*e*_, *F*_*e*_, *H*_*e*_). Then

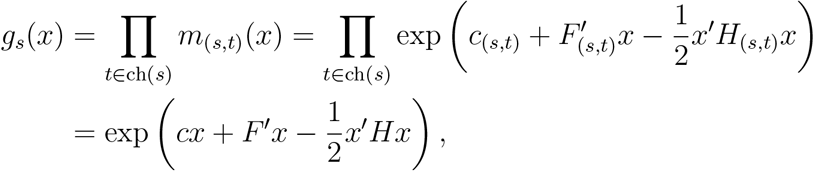

for (*c, F, H*) = ∑_*t*∈ch(*s*)_ *c*_(*s,t*)_, *F*_(*s,t*)_, *H*_(*s,t*)_. Therefore, the fusion step simply consists of adding triplets from all incoming messages. Using rules (1)–(3) will be referred to as the *backward filtering* step. It results in *g* being computed from the leaves back to the root. At any node, or any time-instance on an internal edge, *g* is represented by the triplet (*c, F, H*). We define the *guided process* on the edge *e* by the process *X*^∘^ that solves the SDE

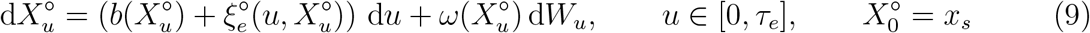

where 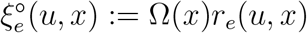 with

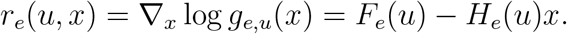

The process *X*^∘^ obtained by simulating this process on the whole tree in a preorder traversal is called the *guided process on the tree*. Denote the laws of *X*^∘^ and *X*^⋆^, considered as process on the whole tree, by ℙ^∘^ and ℙ^⋆^ respectively. It follows from the derivation in Section 4 of (Mider et al., 2021) that

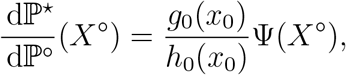

where

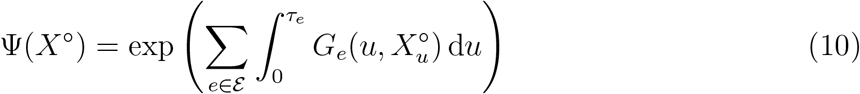

and on edge *e*

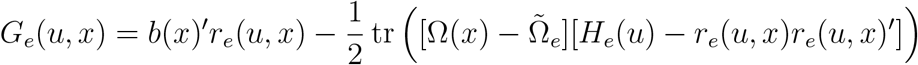

with 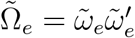. Note that *g*_0_(*x*_0_) is the value of *g* at the root node, evaluated at *x*_0_.

By sampling the process *X*^∘^ we can get weighted samples of the conditioned process. This forms the basis of the MCMC-algorithm that we use. As the backward filtering step is followed by forward simulating the guided process using a preorder traversal, the combined step is called *Backward Filtering Forward Guiding* (BFFG) in Mider et al. (2021).

##### Details on step (2)

We now give details on the second step of the backward filtering. For this, consider the edge *e* = (*s, t*) and suppose we are given 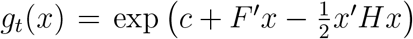 (to reduce notation, we omit dependence on the vertex in denoting the triplet). Examples of choices of 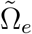 are

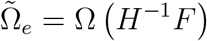

(to be read as Ω(*x*) evaluated at *x* = *H*^−1^*F*), or

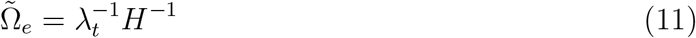

where *λ*_*t*_ *>* 0 at the node *t* at the end of the edge *e* = (*s, t*) is a scalar that can be computed recursively. Specifically *λ*_*t*_ is the observation noise variance when *t* is a leaf node, and 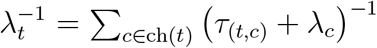 otherwise. The factor has the effect of rescaling the magnitude of *H*^−1^ so that 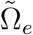 will have consistent magnitude for all edges. In the current implementation, we use this choice. In both cases, solutions to (7) take the form as specified in Equation (8). Moreover,

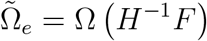

where *u* ∈ [0, *τ*_*e*_] and 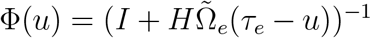. This is a special case of the equations in Theorem 2.5 in Mider et al. (2021) with *B* = 0 and *β* = 0; see also specifically Section 4.2 in Corstanje and van der Meulen (2024)). The computation of *c*_*e*_(*u*) can be omitted as it does not affect the guided process, and it cancels out in the subsequent MCMC acceptance ratios (see (Corstanje and van der Meulen, 2024, Lemma 3.7)).

#### Inclusion of a Super Root

So far, we have assumed *x*_0_ to be known. To address this problem, we define a prior distribution over *x*_0_ by adding an additional node, which we call the super root and denote by *s*, connected to the root of the tree. This super root is assumed to have a known shape, *x*_*s*_. The length of the edge between the root and the super root follows the shape process defined in (3) and has length *l*_*s*_. In this way, the prior distribution on the initial state *x*_0_ is parametrised by (*x*_*s*_, *l*_*s*_). The kernel parameters on the edge connecting the super root and initial state are inferred in the MCMC algorithm and assumed to be the same as the parameters governing the shape process in the rest of the tree.

#### MCMC-algorithms

We introduce an algorithm that samples from the joint distribution of the shape process on the tree and the parameter *θ* = (*α, σ*). The BFFG scheme suggests an MCMC-algorithm where steps 2 and 3 are iterated:

1. *Initialisation*: choose a value for *θ*;
2. *Path imputation*: conditional on *θ*, backward filter and forward guide;
3. *Parameter updating*: conditional on the simulated guided process, update *θ* by a Metropolis-Hastings step.

The difficulty with this scheme is that the resulting Markov chain will be reducible, see Roberts and Stramer (2001). In other words, the chain will not move. Intuitively, once we fix a path of the diffusion process, we can determine the parameters from the quadratic variation of the process. For this reason, we will reparametrise the guided process on each internal edge, as detailed for example in van der Meulen and Schauer (2017). Consider the SDE (9) for the guided process on an edge *e* = (*s, t*). We assume there exists a strong solution. This means that the solution can be constructed from its initial state *x*_*s*_ and the driving Wiener process *W*. From the point of view of implementation this is clear: once we know the starting value *x*_*s*_ and the increments of the Wiener process on a grid, we can discretise the SDE and approximate its solution on that grid. Mathematically, there exists a map ℱ_*θ*_ such that the guided process on the edge *e* is given by ℱ_*θ*_(*x*_*s*_, *W*_*e*_). Define the collection of all driving Wiener processes by *W*, i.e. *W* = (*W*_*e*_, *e* ∈ ℰ). The algorithm we then propose is a Gibbs sampler on (*W, θ*). The reparametrisation in terms of *W* resolves the reducibility problem of the initially proposed algorithm. For updating *W* we use a pre-conditioned Crank-Nicolson step (pCN), cf. Cotter et al. (2013). The overall algorithm is an adaptation of the algorithm proposed in Section 4 of Mider et al. (2021), where we adapt from a hidden Markov model to a directed tree and take the specific structure in (3) into account. We note that in our implementation the parameters are updated componentwise, i.e., individually.

#### Prior and Proposal Specification

In this section we describe the details of the MCMC algorithm. The specific choice of parameters for a given experiment is described in the section *“Results”*. With slight abuse of notation, we will write *X*^∘^ = ℱ_*θ*_(*x*_*s*_, *W*) to denote the guided process on the tree (rather than on a single edge). Recall the definition of Ψ(*X*) in (10). To highlight the dependence of Ψ, as also *g*, on *θ* we will add “; *θ*” to the algorithmic description of the algorithm.

##### Prior distributions

- The prior on the root node is defined using (*x*_*s*_, *l*_*s*_) where *x*_*s*_ is the pre-selected shape at the super root and *l*_*s*_ is the length of the branch going from the super root to the root.
- The prior on the kernel parameters is uniform in a fixed interval, i.e., *θ* = (*α, σ*) ~ 𝒰 (*a*_*α*_, *b*_*α*_) ⊗ 𝒰 (*a*_*σ*_, *b*_*σ*_). We denote this density by *π*.

##### Metropolis-Hastings proposals

- pCN: conditional on *W*, on each edge *e*, we propose

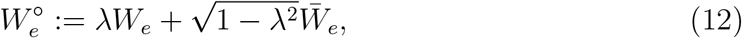

where 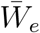 is a Wiener process on edge *e* that is independent of *W*_*e*_. In our MCMC algorithm we need to specify the correlation parameter *λ* ∈ [0, 1).
- Given the boundaries *a* and *b* where *a < b*, along with standard deviation *τ*, we propose a value for *κ*^∘^ conditional on the current value of the parameter *κ* by sampling from the symmetric reflected Gaussian distribution, *q*(· | *κ, τ*), as described in Appendix 2.

##### Other parameters

- *N*, the number of MCMC iterations.
- *δ*, the stepsize or mesh-width for numerically approximating the SDE for the guided process.
- *γ*, the variance of the extrinsic noise (i.e. the observational noise) which we assume to follow a multivariate Gaussian distribution where the covariance matrix is given by *γ* · *I*.

###### Algorithm 1

Metropolis-Hastings algorithm

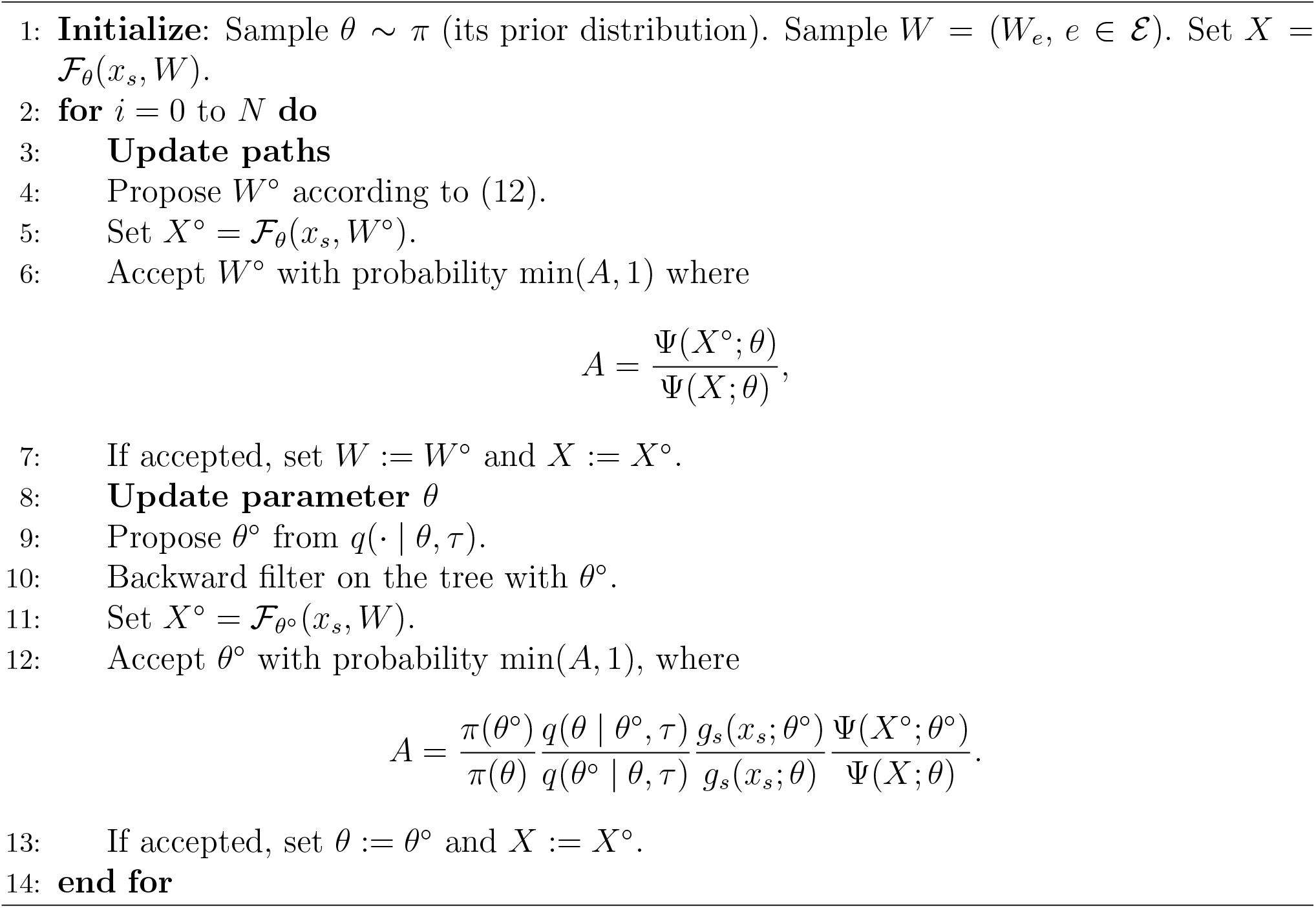

### Simulation Studies

In this paper, we conduct five simulation studies, each described in detail in this section. We start by describing settings which are shared across studies and following this, we describe the specifics of each simulation study.

For all simulation studies, datasets are simulated on different topologies and with different parameters using the stochastic shape model introduced in section *“Model and Parameters”*. For all simulated datasets a stepsize of *δ* = 0.05 is used for discretizing the stochastic differential equation that captures shape evolution. Shapes used as root for simulation include either a downsampled *Morpho hercules* forewing from Chazot et al. (2021) (which we will refer to as butterfly forewing) or a circle (see Fig. 3). Both shapes are made up of 20 landmarks in 2D. The number of noise fields *J* is set to match the number of landmarks, and the diffusion matrix corresponds to the noise fields being at the landmark positions. Phylogenies used for simulation include the full phylogeny from Chazot et al. (2021) (see Fig. 7), a subtree of the full phylogeny with or without a root branch (see Fig. 4a and 4b), a symmetric phylogeny (see Fig. 4c), and an asymmetric phylogeny (see Fig. 4d).

**Figure 3.**
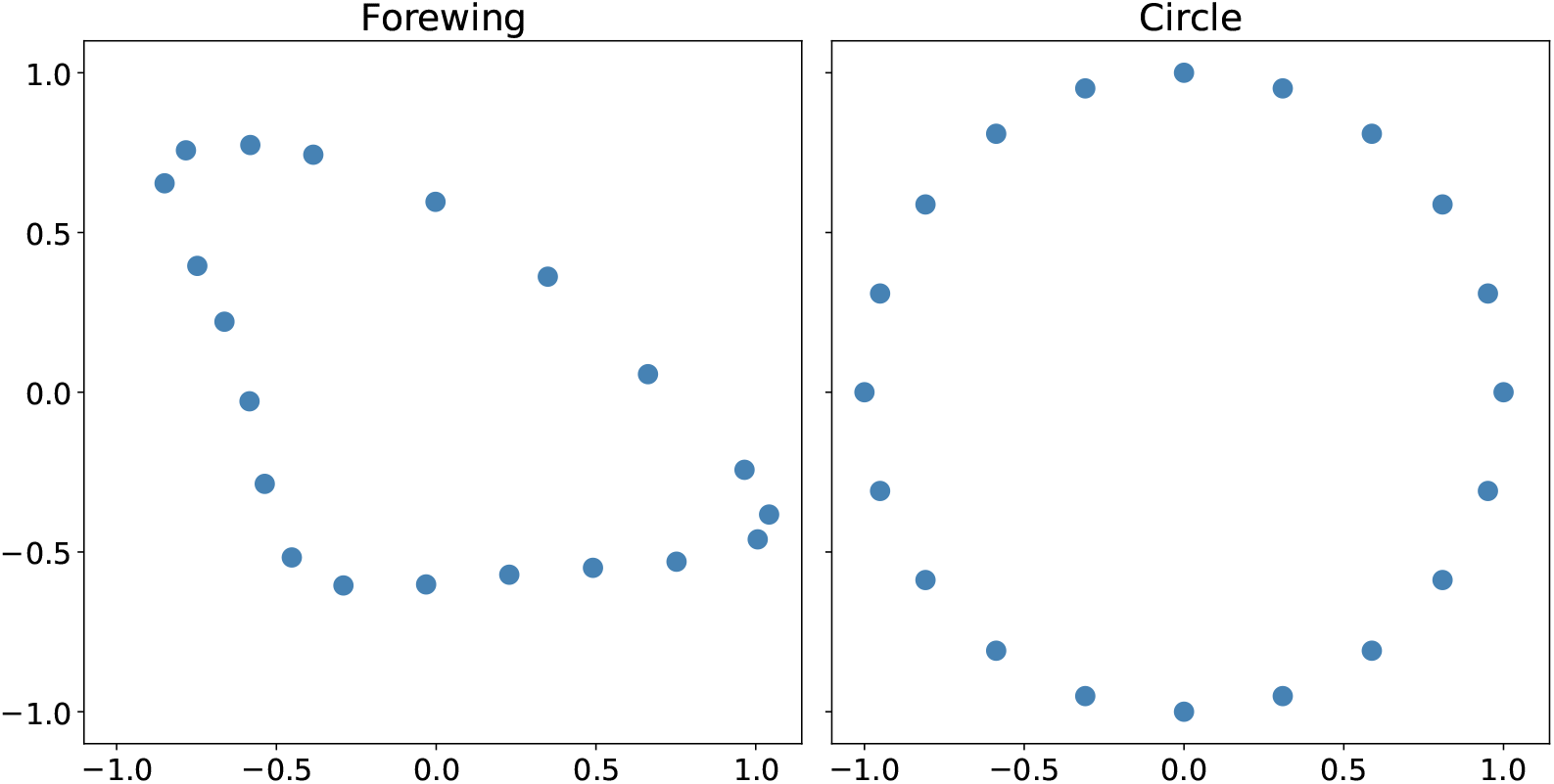
Left: downsampled *Morpho hercules* forewing. Right: circle.

**Figure 4.**
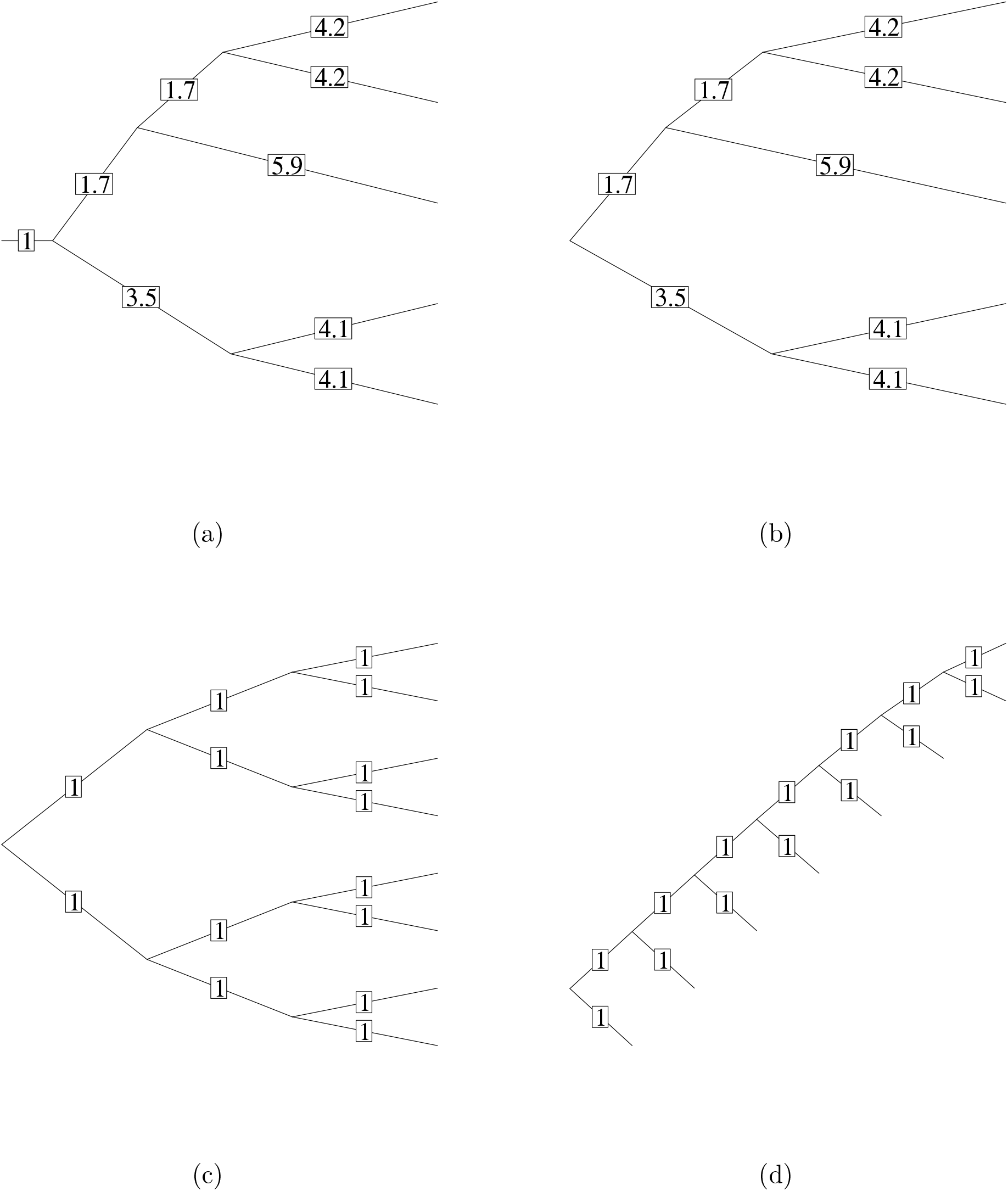
Phylogenies used for simulation studies. (a) Mixed phylogeny with a root branch of length 1. (b) Mixed phylogeny. (c) Symmetric phylogeny. (d) Asymmetric phylogeny.

In all simulation studies which include estimation of a posterior, we use the same stepsize as we use to simulate the data, i.e., *δ* = 0.05. Each posterior distribution is inferred using three MCMC chains initialized by values sampled from the prior and subsequent samples from the guided process. We assess convergence for each of the inferred posterior using standard convergence diagnostics in form of an improved version of the Gelman-Rubin statistic 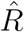 (Vehtari et al., 2021) as implemented in the Python package ArviZ (Kumar et al., 2019). In the experiments where we estimate a posterior mode, we do this marginally using the kernel density estimator from ArviZ. Bandwidth is estimated using the Silverman estimator.

#### Simulation Study 1: Evaluation of Metropolis-Hastings Algorithm

In this simulation study, the aim is to validate the implementation of our proposed Metropolis-Hastings algorithm. For this we use Simulation-Based Calibration (SBC) proposed by Talts et al. (2018). The idea underlying SBC is that for a Bayesian model the data averaged posterior reduces to the prior distribution when the data is simulated from the model. This implies that the order statistic of the true value relative to the posterior sample should be uniformly distributed, and so we can use this property to assess whether a Bayesian method has been correctly implemented. We note that this type of assessment is a stronger requirement than the often used MCMC check involving convergence of the posterior to the prior when running the method without data, as the evaluation method also require the likelihood calculations to be implemented correctly. It ensures that the posterior probabilities are calibrated correctly. Lastly, we also note that due to the difficulty of defining a prior distribution for shapes, we are only able to validate the Metropolis-Hastings algorithm using SBC conditioned on the true root. For further explanation on SBC we refer to Appendix 3.

In order to run SBC we simulate 100 datasets on the phylogeny given in 4a with the root shape being the butterfly forewing shown in Figure 3. Each of the 100 datasets are simulated using a different set of parameters sampled from the prior distributions *α* ~ 𝒰 (0.0005, 0.03) and *σ* ~ 𝒰 (0.7, 1.3). The 100 combinations of parameters sampled from the prior and used for simulation of the datasets are available as Supplementary Material (Fig. S3). In order to simulate from the full probabilistic model (including measurement noise), isotropic Gaussian noise with a variance of *γ* = 0.001 is added to the simulated shapes. We infer the posterior by running Algorithm 1 with the following settings; *λ* = 0.8, *τ*_*α*_ = 0.007, *τ*_*σ*_ = 0.2, and *γ* = 0.001. In order to run SBC we let the root be equal to the true root, i.e. the butterfly forewing given in Figure 3.

Finally, for readers interested in the frequentist properties of the estimators, we also evaluate the bias for the posterior mean, the posterior median and the posterior mode. For each of the 100 simulated data sets we compute the difference between the true value of the parameter, *θ*, and the posterior point estimate, 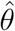, i.e. 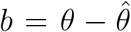. We estimate bias as the empirical mean of *b* i.e. 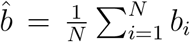. As the kernel parameters have very different range, we divide *b* with the mean of the true parameter i.e. 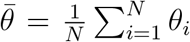 where *θ*_*i*_ refers to the parameter used for simulation of each of the *i* data sets.

#### Simulation Study 2: Evaluation of Root Estimation Strategy

In this simulation study, the aim is to evaluate our root estimation strategy. We do this by comparing the MSE for root estimates obtained using our method and root estimates obtained using standard methods. We simulate data on three different phylogenies and with two different roots. Phylogenies used are given in Figure 4b, c, and d. Root shapes used are shown in Figure 3. Datasets are simulated for each combination of phylogeny, parameters and root as shown in Table 1. We pick kernel parameters such that the datasets contain a reasonable amount of variation. In order to simulate from the full probabilistic model, isotropic Gaussian noise with a variance of *γ* = 0.001 is added to the simulated shapes. Visualization of all simulated data sets is available as Supplementary Material (Fig. S14, S15, S16, S17, and S18).

**Table 1.**
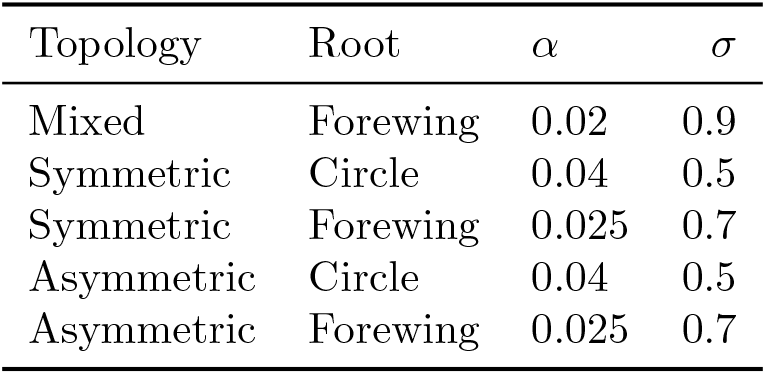
Table showing different combinations of parameters, phylogeny, and root used for simulating data sets.

For each simulated data set we infer the posterior by running Algorithm 1. For all topologies, root, and parameter combinations we use *γ* = 0.001 and *l*_*s*_ = 2 in our MCMC (see Supplementary Material Fig. S34, S35, and S36 for a visualization of the phylogenies with super root branch). In addition, we let the super root *x*_*s*_ for all settings be the phylogenetic mean of the data. We compute the phylogenetic mean as described in equation (3) in Revell (2009). MCMC settings used for inferring the posterior for each of the simulated data sets is shown in Table 2.

**Table 2.** Table showing the MCMC settings used for inferring the posterior distribution.

| Topology | Root | prior $\alpha$ | prior $\sigma$ | MCMC settings |
| --- | --- | --- | --- | --- |
| Mixed | Forewing | $\mathcal{U}(0.005, 0.03)$ | $\mathcal{U}(0.7, 1.3)$ | $\lambda = 0.8, \tau_\alpha = 0.005, \tau_\sigma = 0.2, N = 3000$ |
| Symmetric | Circle | $\mathcal{U}(0.0, 0.05)$ | $\mathcal{U}(0.0, 1.5)$ | $\lambda = 0.8, \tau_\alpha = 0.005, \tau_\sigma = 0.2, N = 4000$ |
| Symmetric | Forewing | $\mathcal{U}(0.0, 0.05)$ | $\mathcal{U}(0.0, 1.5)$ | $\lambda = 0.8, \tau_\alpha = 0.005, \tau_\sigma = 0.2, N = 4000$ |
| Asymmetric | Circle | $\mathcal{U}(0.0, 0.05)$ | $\mathcal{U}(0.0, 1.5)$ | $\lambda = 0.95, \tau_\alpha = 0.005, \tau_\sigma = 0.05, N = 4000$ |
| Asymmetric | Forewing | $\mathcal{U}(0.0, 0.05)$ | $\mathcal{U}(0.0, 1.5)$ | $\lambda = 0.85, \tau_\alpha = 0.003, \tau_\sigma = 0.2, N = 4000$ |

#### Simulation Study 3: Probabilistic Ancestral State Reconstruction

In this simulation study, the aim is to illustrate our method by carrying out a simulation study on the full phylogeny from Chazot et al. (2021) (see Fig. 7). The root used for simulation is the butterfly forewing shown in Figure 3. The parameters used for simulation are *α* = 0.01 and *σ* = 0.6. We infer the posterior distribution using Algorithm 1 with *l*_*s*_ = 5, *γ* = 0.002, and the super root (*x*_*s*_) being the phylogenetic mean of the observed data. We assume the following priors over the parameters: *α* ~ 𝒰 (0, 0.03) and *σ* ~ 𝒰 (0.4, 1.0). In addition we run the MCMC algorithm for 6000 iterations with a Crank-Nicholson *λ* = 0.97 and with proposal parameters *τ*_*α*_ = 0.003 and *τ*_*σ*_ = 0.07.

#### Simulation Study 4: Comparison of Uncertainty Estimates

In this simulation study, the aim is to compare ancestral state uncertainty estimates obtained from our method with uncertainty estimates obtained from traditional non-shape aware models. In particular, we compare the confidence ellipses obtained for an ancestral state estimate under a Brownian motion model to the posterior distribution obtained using our model.

We simulate data on the phylogeny given in Figure 4b and let the root shape be the butterfly forewing (see Fig. 3). The parameters used for simulation are *α* = 0.03 and *σ* = 0.4. Following simulation, data are Procrustes aligned and rotated in order to be correctly oriented in the coordinate system (this is only done for visualization purposes). We Procrustes align the data using the *gpagen* function in the R package geomorph (Baken et al., 2021; Adams et al., 2025).

We infer the posterior distribution over the ancestral states as well as parameters by running our Metropolis-Hastings algorithm with the following settings; *N* = 3000, *λ* = 0.95, *l*_*s*_ = 1, *γ* = 0.0001, *τ*_*α*_ = 0.005, and *τ*_*σ*_ = 0.1. We assume the following priors over the parameters *α* ~ 𝒰 (0.0, 0.04) and *σ* ~ 𝒰 (0.0, 1.0).

We do maximum likelihood based ancestral state reconstruction under a Brownian motion model and obtain variances for the estimates using the *fastAnc* function in the R package phytools (Revell, 2024). As the likelihood is Gaussian, we show the uncertainty of the ancestral state estimates as confidence ellipses. We estimate confidence ellipses based on the variance estimates obtained for each coordinate.

#### Simulation Study 5: Comparison of Integration Measures

In this simulation study, the aim is to relate the *σ* parameter in our shape model to an existing metric for biological integration. Traditionally, biological integration has been evaluated using eigenvalue dispersion indices. In this simulation study we use a specific type of eigenvalue dispersion index called the relative eigenvalue variance, *V*_*rel*_ Pavlicev et al. (2009) for evaluating the strength of integration for a simulated data set. In our experiments we compute *V*_*rel*_ from the trait covariance matrix as described in Watanabe (2022) and Conaway and Adams (2022). Specifically, we use the *integration*.*Vrel* function from geomorph (Baken et al., 2021; Adams et al., 2025) to compute *V*_*rel*_ from the non-trivial dimensions of variance (Conaway and Adams, 2022).

We simulate datasets on a star phylogeny with 100 tips and where all branches have length 1. For each value of *σ* we simulate 50 data sets using *α* = 0.05, and with the root shape being the butterfly forewing shown in Figure 3. We compute *V*_*rel*_ for the same data sets with or without Procrustes alignment.

### Empirical Data Set

In section *“Analysis of Morpho Butterflies”* we analyze the butterfly data set from Chazot et al. (2021) containing phylogeny and landmark data of the *Morpho* genus. Specifically, we analyze the landmarked shapes of the forewing of male butterflies. In case several specimens have been landmarked we used the euclidean average of all the specimen as our data. In the original data set, each forewing was landmarked with 44 landmarks of which 18 were normal landmarks and 26 were semilandmarks. In order to increase computational speed, we downsampled the number of landmarks to 20 keeping only landmarks denoting the outline of the forewing. Procrustes alignment of the forewing was made based on the full data set of 44 landmarks.

We infer the posterior over parameters and paths using Algorithm 1 with *l*_*s*_ = 5 and the super root (*x*_*s*_) being the phylogenetic mean of the observed data. We assume the following priors over the parameters: *α* ~ 𝒰 (0.0, 0.01) and *σ* ~ 𝒰 (0.0, 1.5). In addition, we run the MCMC algorithm for 10, 000 iterations with a Crank-Nicholson *λ* = 0.99, observation noise *γ* = 0.001, stepsize *δ* = 0.05, and with proposal parameters *τ*_*α*_ = 0.0015 and *τ*_*σ*_ = 0.1. We assess convergence for each of the inferred posterior using standard convergence diagnostics in form of an improved version of the Gelman-Rubin statistic 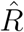 (Vehtari et al., 2021) as implemented in the Python package ArviZ (Kumar et al., 2019). The posterior distribution is inferred using three MCMC chains initialized by values sampled from the prior and subsequent samples from the guided process. We estimate the posterior mode marginally using the kernel density estimator from ArviZ. Bandwidth is estimated using the Silverman estimator.

## Results

### MCMC Algorithm Yields Well Calibrated Credible Sets

In Figure 5, we show trace plots and a marginal posterior densities obtained for a single data set for the parameter *σ* and the parameter *α*. Generally, we obtain 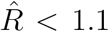 indicating convergence. Distributions of 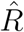 for all dimensions of all internal nodes as well as parameters are available as Supplementary Material (Fig. S4, S5, S6, S7, and S8). For 83 out of 100 data sets convergence was achieved with standard settings. For the remaining 17 data sets convergence required more correlated proposals and/or more iterations. Specifically, increasing the number of MCMC iterations from 3000 to 5000 alone led to convergence for one data set, and additionally increasing the correlation between proposed paths i.e. *λ* from 0.8 to 0.85 (3 data sets), 0.9 (2 data sets), and 0.95 (9 data sets) resulted in convergence for most data sets. For the remaining two data sets convergence was achieved by increasing the MCMC iterations to 7000 and increasing the correlation between proposed paths to *λ* = 0.97. For one of the two data sets we also increased the correlation between the proposed *σ* parameter by decreasing the proposal standard deviation i.e. setting *τ*_*σ*_ = 0.1. We note that generally achieving MCMC convergence with our default settings was particularly challenging when data sets were simulated with large *α* and small *σ*. The distribution of rank statistics for all dimensions of all landmarks in all internal nodes and parameters is available as Supplementary Material (Fig. S9, S10, S11, S12, and S13). In addition to visually inspecting the rank statistics distributions we also carry out a Kolmogorov-Smirnov test on the rank distribution obtained for each dimension of the posterior to test for uniformity. We find that 1 out of the 162 tests is below our significance level of 1%. Given that we carry out 162 tests this is close to the number of significant test statistics we would expect. This suggests that the rank statistics are indeed uniformly distributed and that our Metropolis-Hastings algorithm is implemented correctly.

**Figure 5.**
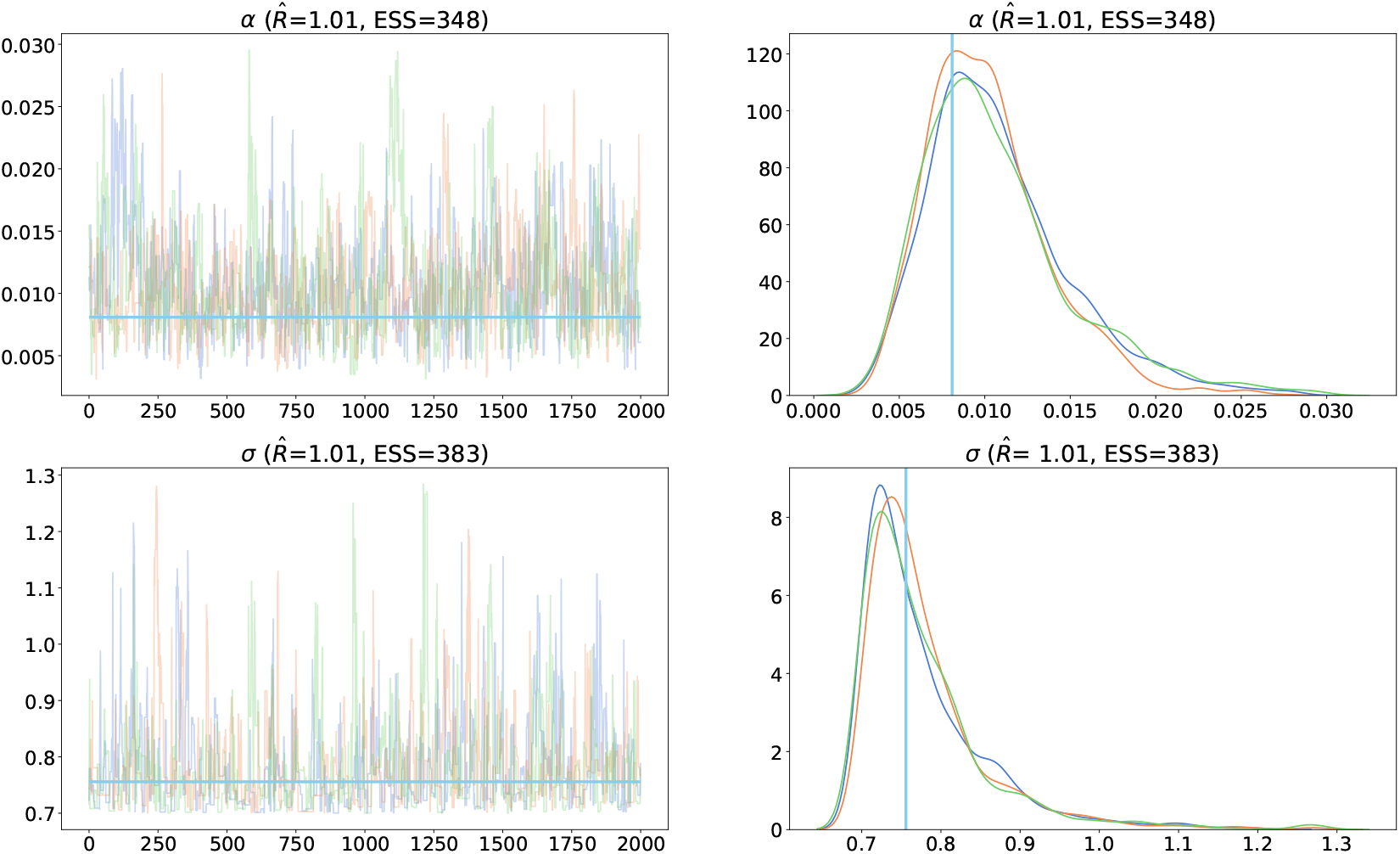
Trace plots and marginal posterior densities obtained for a single data set. Colors indicate the result for 3 independent chains. Trace plots and density plots are shown after removal of burnin (first 1000 iterations). Leftmost column shows trace plots obtained for a single data set for the parameter *α* (upper row) and the parameter *σ* (lower row). Rightmost column shows the corresponding marginal posterior density. 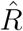 refer to an improved version of the Gelman-Rubin convergence diagnostics (Vehtari et al., 2021). ESS is the effective sample size.

In addition to running SBC we also investigate the bias of the posterior point estimates. We show the distribution of 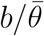 in Figure 6. We compute standard 99% Waldtype confidence for the bias estimate 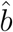 for each of the estimators. For the kernel parameter *σ* we obtain the following confidence intervals; 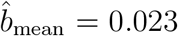 (99% CI [−0.008, 0.054]), 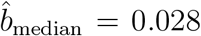 (99% CI [−0.003, 0.059]) and 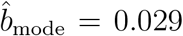 (99% CI [−0.015, 0.073]). For the kernel parameter *α* we obtain the following confidence intervals; 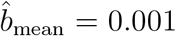 (99% CI [−1.653 · 10^−5^, 0.002]), 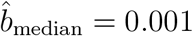 (99% CI [4.054 · 10^−4^, 0.002]) and 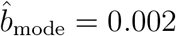 (99% CI [0.001, 0.003]). Based on the confidence intervals we conclude that all posterior estimators of the parameter *σ* are unbiased, whereas only the mean estimator for the parameter *α* is unbiased. We note that Bayesian estimators are not guaranteed to be unbiased.

**Figure 6.**
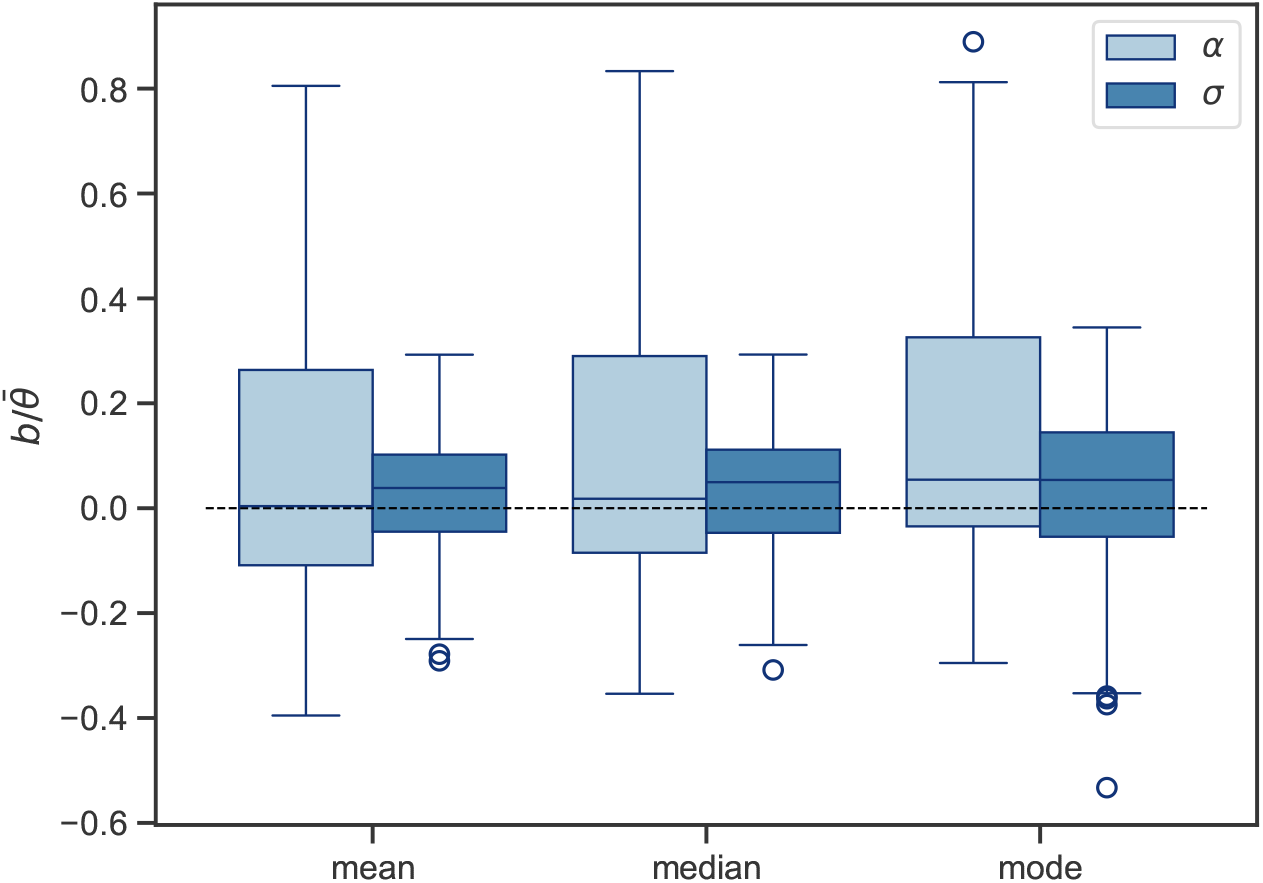
Figure showing boxplots of 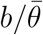 for all posterior point estimators. The mode of the marginal distribution of either *α* or *σ* is estimated using the kernel density estimator implemented in the Python package ArviZ (Kumar et al., 2019) with bandwidth selected using the Silverman estimator.

### Root Estimation

For all data sets simulated on a mixed topology with the root shape being a butterfly forewing we obtain 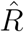 values close to 1 suggesting that the chains have converged (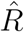 for all runs are available as Supplementary Material Fig. S19). All data sets simulated on a symmetric topology with the root shape being a butterfly forewing achieved approximate convergence with the settings described in Table 2. For the data set simulated on a symmetric topology with the root shape being a circle, 18 out 20 data sets achieved approximate convergence with the MCMC settings shown in Table 2. We achieved approximate convergence for the last two data sets by increasing the number of MCMC iterations to 6000, increasing the correlation between proposed values of *α*, i.e., setting *τ*_*α*_ = 0.002 and increasing the Crank-Nicholson correlation parameter *λ*, i.e., setting *λ* = 0.9. Convergence diagnostics for all estimated posterior distributions for data sets simulated on symmetric topologies are available as Supplementary Material (Fig. S20 and S21). All data sets simulated on an asymmetric topology with the root shape being a circle achieved approximmate convergence with the MCMC settings described in Table 2. For the data sets simulated on an asymmetric topology with the root shape being a butterfly forewing, 17 out of 20 data sets achieved approximate convergence using the MCMC settings described in Table 2. For the remaining three data sets approximate convergence was achieved by setting *N* = 6000 and *λ* = 0.9 (one data set), setting *N* = 6000 and *λ* = 0.95 (one data set), or setting *N* = 10000 and *λ* = 0.99 (one data set). Convergence diagnostics for all estimated posterior distributions for asymmetric topology are available as Supplementary Material (Fig. S22 and S23).

We compute the Mean Squared Error (MSE) between a selected point estimate and the true root and compare the MSE obtained for different posterior point estimates with standard point estimates, i.e., the mean and the phylogenetic mean of the data. In Table 3 we show the MSE for different point estimates of the root. As can be seen in Table 3 we obtain the smallest MSE with posterior point estimates for all combinations of topologies, root shapes, and parameters. This suggests that the proposed Bayesian approach in fact does succeed in improving on the phylogenetic mean estimate. We note that the MSE for the posterior median is lower than the mean and phylogenetic mean in all experiments. Therefore, we recommend using the posterior median for practical data analysis.

**Table 3.** MSE for different point estimates of the root. MSE is computed based on 20 simulated data sets for all topologies and root combinations. The mode is estimated marginally using the kernel density estimator implemented in the Python package ArviZ (Kumar et al., 2019) with bandwidth selected using the Silverman estimator. Data sets were simulated with the parameters stated in Table 1. For each experiment we show the smallest MSE in bold.

| Topology | Root | Classical point estimators |  | Posterior point estimators |  |  |
| --- | --- | --- | --- | --- | --- | --- |
|  |  | Mean | Phylogenetic mean | Mean | Median | Mode |
| Mixed | Forewing | $5.967 \cdot 10^{-3}$ | $5.761 \cdot 10^{-3}$ | **$5.651 \cdot 10^{-3}$** | $5.664 \cdot 10^{-3}$ | $5.719 \cdot 10^{-3}$ |
| Symmetric | Circle | $4.889 \cdot 10^{-3}$ | $4.889 \cdot 10^{-3}$ | **$4.753 \cdot 10^{-3}$** | $4.763 \cdot 10^{-3}$ | $4.889 \cdot 10^{-3}$ |
| Symmetric | Forewing | $3.153 \cdot 10^{-3}$ | $3.153 \cdot 10^{-3}$ | $3.164 \cdot 10^{-3}$ | **$3.138 \cdot 10^{-3}$** | $3.148 \cdot 10^{-3}$ |
| Asymmetric | Circle | $9.621 \cdot 10^{-3}$ | $3.395 \cdot 10^{-3}$ | $3.348 \cdot 10^{-3}$ | $3.341 \cdot 10^{-3}$ | **$3.328 \cdot 10^{-3}$** |
| Asymmetric | Forewing | $7.700 \cdot 10^{-3}$ | $2.481 \cdot 10^{-3}$ | $2.421 \cdot 10^{-3}$ | **$2.416 \cdot 10^{-3}$** | $2.474 \cdot 10^{-3}$ |

### Probabilistic Ancestral Reconstruction

For almost all dimensions of all internal nodes the 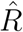-statistic is below 1.1 suggesting that the MCMC chains have converged. The 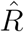-statistic for all internal nodes is available as Supplementary Material (Fig. S24). In Figure 7 we show the posterior obtained for all internal nodes. As can be seen, the posterior recovers the true shape quite well in all internal nodes. From Figure 7 we also see that the uncertainty differs depending on the position in the tree. Specifically, we note that the shapes which are estimated with the smallest uncertainty (i.e. node 1, 34, and 36) are the nodes that are closest to the observed leaves. This observation is not unexpected as the model has more information about the shape in nodes close to leaf nodes that harbor the observed data.

**Figure 7.**
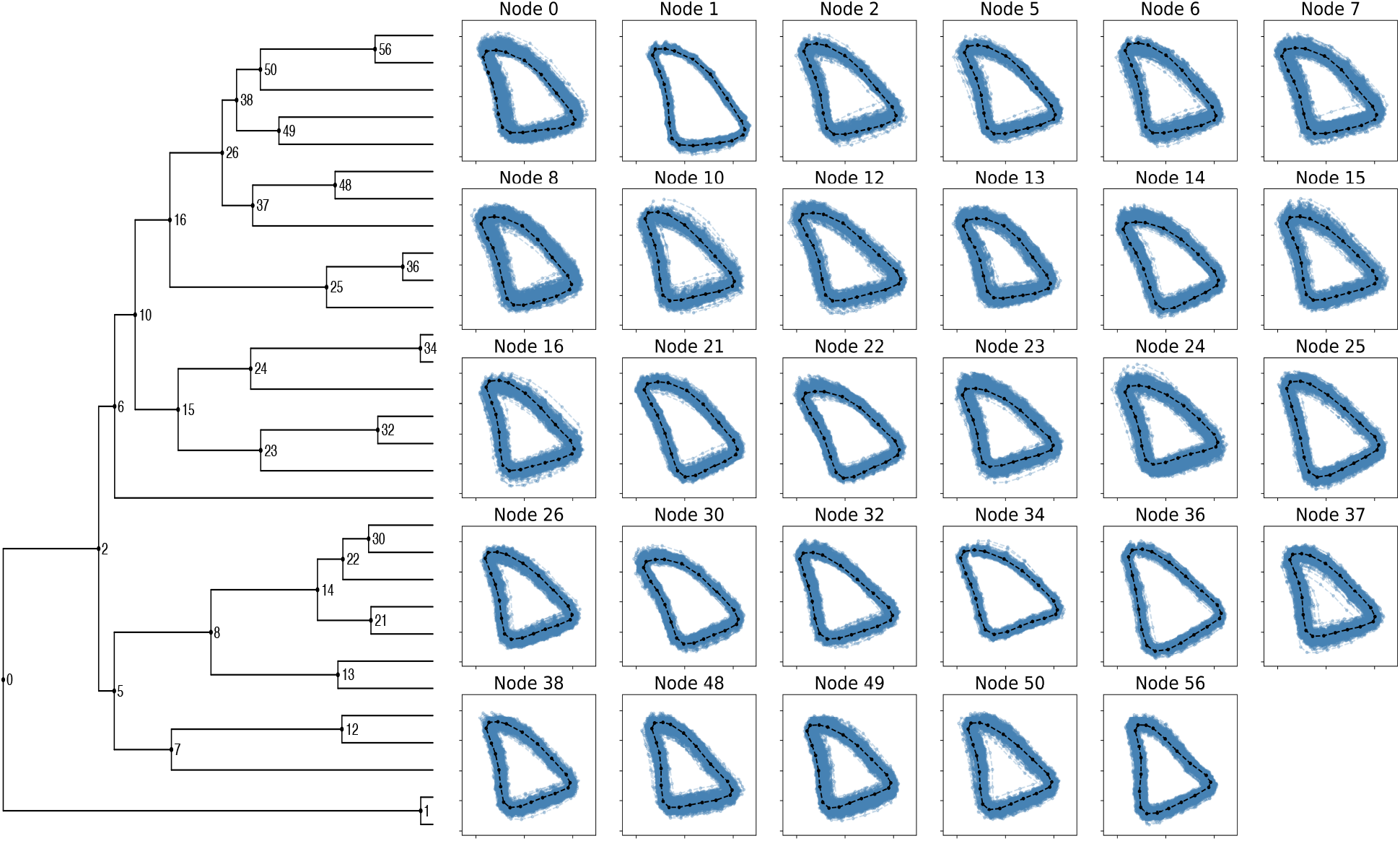
Figure showing the posterior distribution over the internal nodes. The phylogeny to the left is the phylogeny from Chazot et al. (2021) which we use for simulating the data set. To the right we show samples from the posterior distribution (given in blue). The shape given in black is the true shape at the specific node. 6000 MCMC iterations were used for running the chains and a burnin of 2000 iterations has been removed prior to making these plots.

In Figure 8 we show trace plots and density plots for the posterior obtained for the kernel parameters *α* and *σ*. Based on the 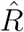 values and the trace plots we conclude that the chains have approximately converged. For the *α* parameter we estimate the posterior mean to be 0.013 and a 95% credible interval (i.e. highest posterior density interval) of (0.008, 0.017). For *σ* we estimate the posterior mean to be 0.728 with credible interval (0.598, 0.849). As we have simulated the data, we know the true values of the kernel parameters (*α* = 0.01 and *σ* = 0.6). These true values lie within the credible interval.

**Figure 8.**
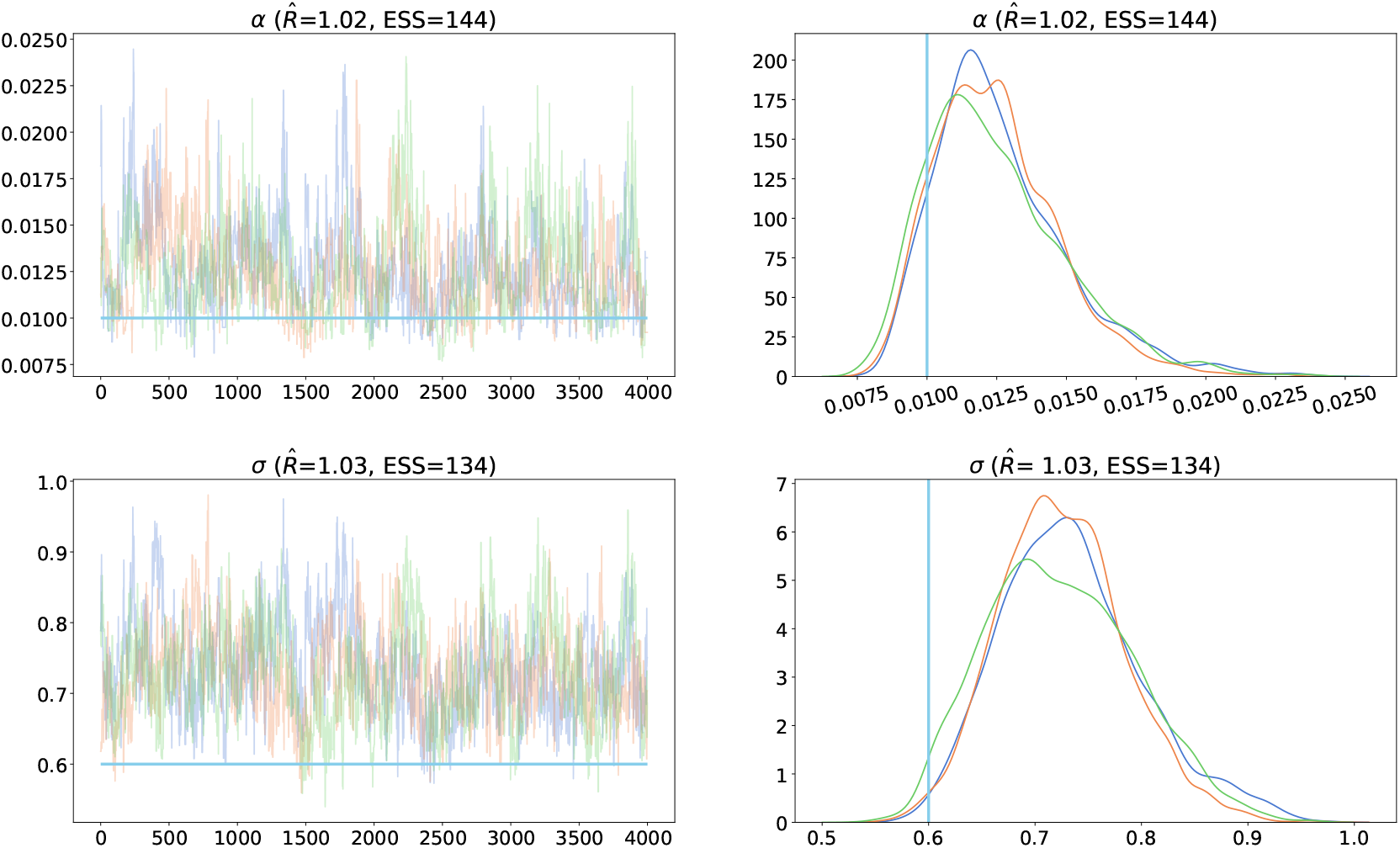
Figure showing the posterior distribution obtained for the different kernel parameters *α* and *σ*. Colors indicate the result for 3 independent chains. Leftmost column shows trace plots and rightmost column shows the marginal posterior densities for the different MCMC chains. Trace plots and density plots are shown after removal of burnin (first 2000 iterations) 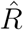 refers to an improved version of the Gelman-Rubin convergence diagnostics (Vehtari et al., 2021). ESS is the effective sample size.

### Shape-Aware Model Yield Meaningful Uncertainty Estimates

We assess convergence using standard convergence diagnostics and conclude that our MCMC chains have converged (see Supplementary Material Fig. S26, S27, S28, S29, S30 for traces and 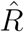 diagnostics). In Figure 9 we show confidence ellipses for each landmark in the estimated ancestral state and samples from the posterior distribution. As can be seen in Figure 9, the confidence ellipses computed using traditional methods allow landmarks to overlap. This is different from uncertainty estimates obtained using our model, as our uncertainty bounds are obtained from a (joint) posterior distribution over all landmarks and based on a shape aware stochastic model which does not allow landmarks to overlap. As landmarks typically represent the outline of a physical object we would not expect any landmarks in our ancestral shapes to overlap. This emphasizes the need for joint modeling of all landmarks in order to achieve meaningful uncertainty bounds on the ancestral reconstructions.

**Figure 9.**
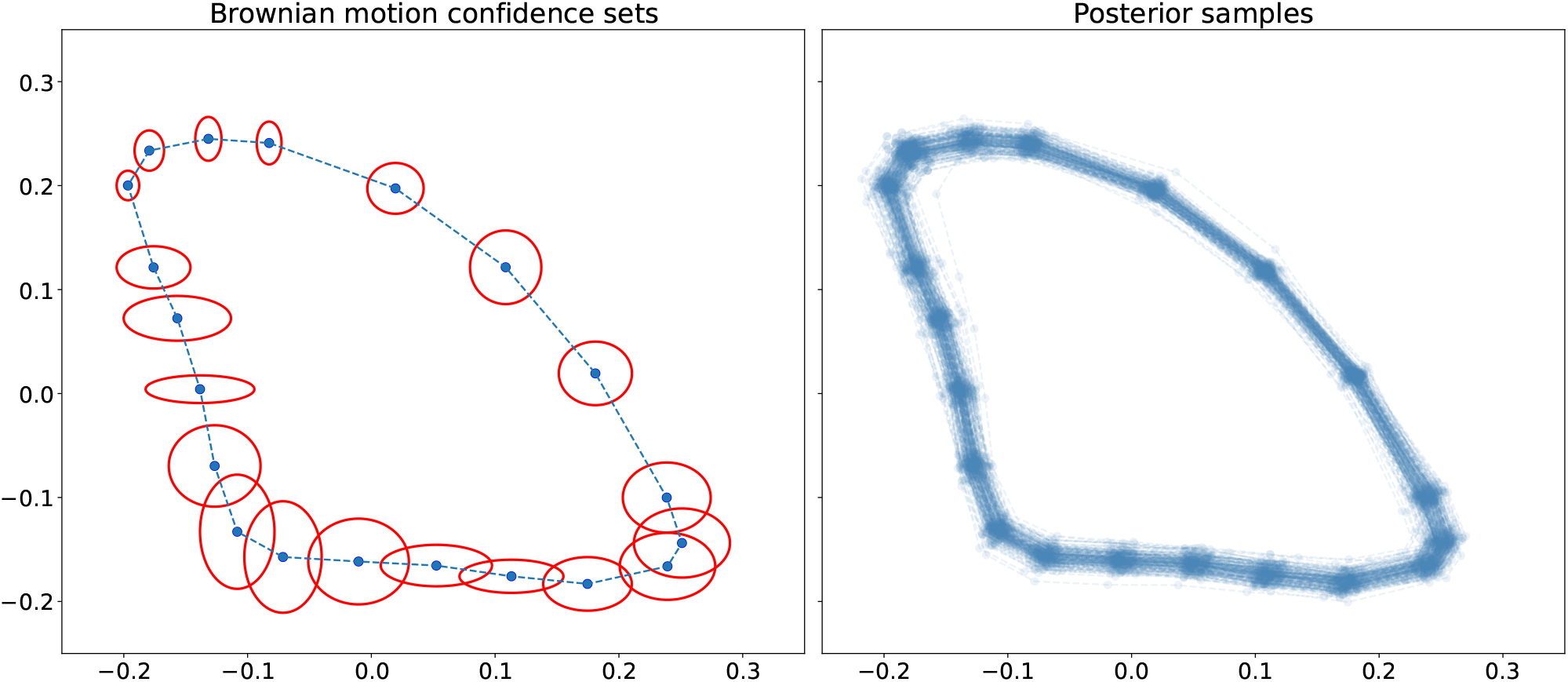
Figure showing uncertainty estimates for a reconstructed shape. Left: 95% confidence ellipse for a ancestral reconstructed shape inferred using the *fastAnc* function from the R package phytools. Right: posterior samples for the reconstructed shapes. We plot every 100 posterior sample from the MCMC chains after removing a burnin of 30%.

### *σ* Parameter as a Measure of Biological Integration

In Figure 10 we show *V*_*rel*_ for the 50 different data sets simulated for each value of *σ*. As can be seen in Figure 10 the integration measure, *V*_*rel*_ and the *σ* parameter are strongly positively correlated, which suggests that the *σ* parameter quantifies correlation in shape in a manner very similar to the standard measure of biological integration. We also note that, when using Procrustes alignment, there is a saturation effect, where for large values of *σ*, increases in *σ* are associated with diminishing increases in *V*_*rel*_, suggesting that Procrustes alignment may bias estimation of biological integration. This finding is in line with existing findings, see for example Rohlf and Slice (1990); Klingenberg (2009); Cardini et al. (2019), and Zelditch and Swiderski (2023).

**Figure 10.**
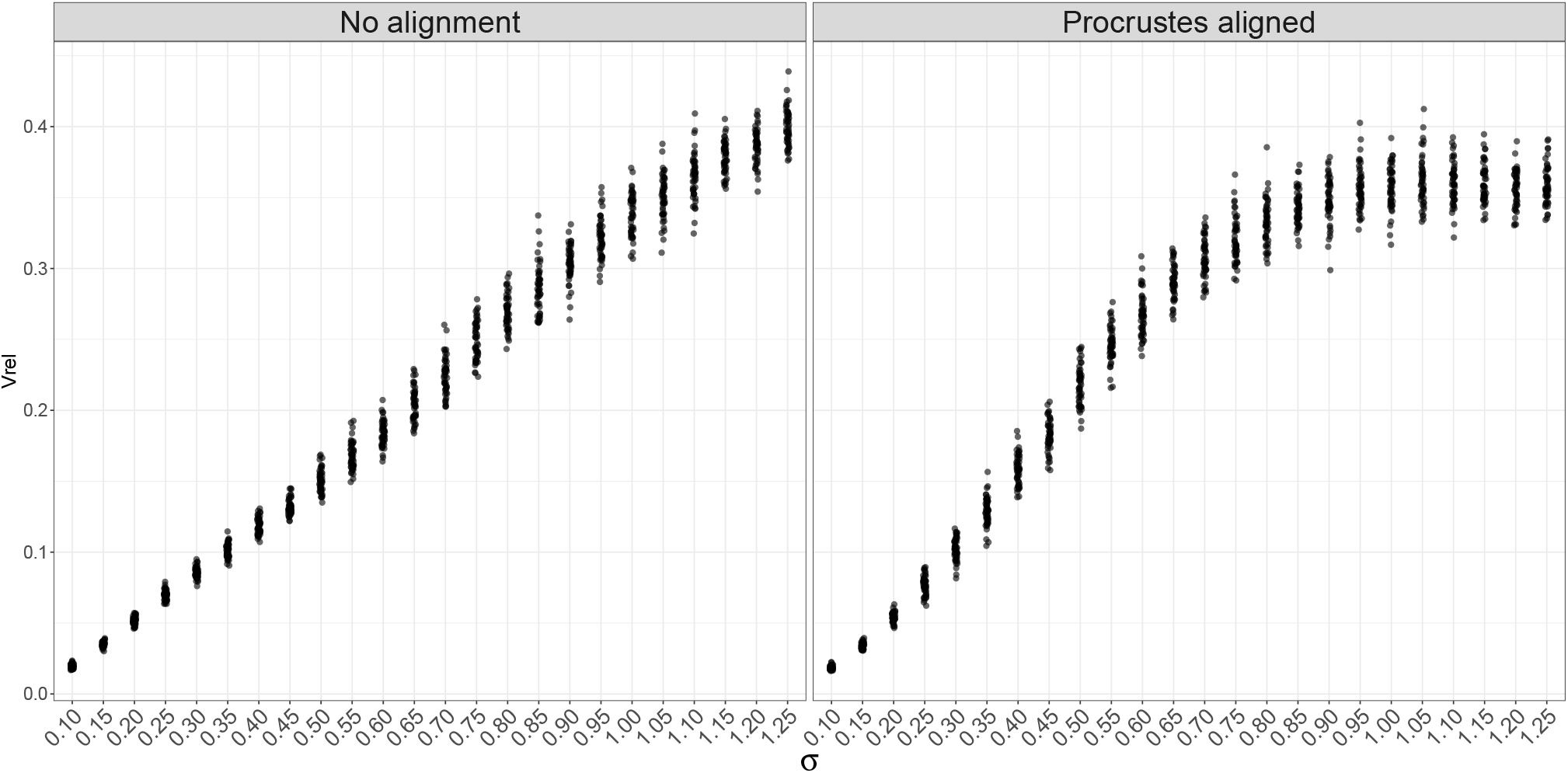
Figure showing *V*_*rel*_ obtained for data sets simulated using different *σ* parameters. Left: *V*_*rel*_ obtained for simulated data that have not been Procrustes aligned. Right: *V*_*rel*_ obtained for simulated data that have been Procrustes aligned.

### Analysis of *Morpho* Butterflies

For almost all dimensions of all internal nodes, the 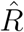-statistic is below 1.1 suggesting that our MCMC chains have approximately converged. The 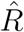-statistic for all internal nodes is available as Supplementary Material (Figure S25). In Figure 11 we show the posterior distribution over all the internal nodes. Although the posterior shapes overall look quite similar we still observe some variation in posterior shapes. For instance, the posterior shape we obtain for node 12 seem to be quite elongated, with a pointy apex (the upper corner of the wing furthest away from the body of the butterfly) and bended termen (the edge of the wings furthest away from the body of the butterfly). On the other hand we also observe posterior shapes that are less elongated and have less bended termen such as the posterior shape which we obtain for node 34.

**Figure 11.**
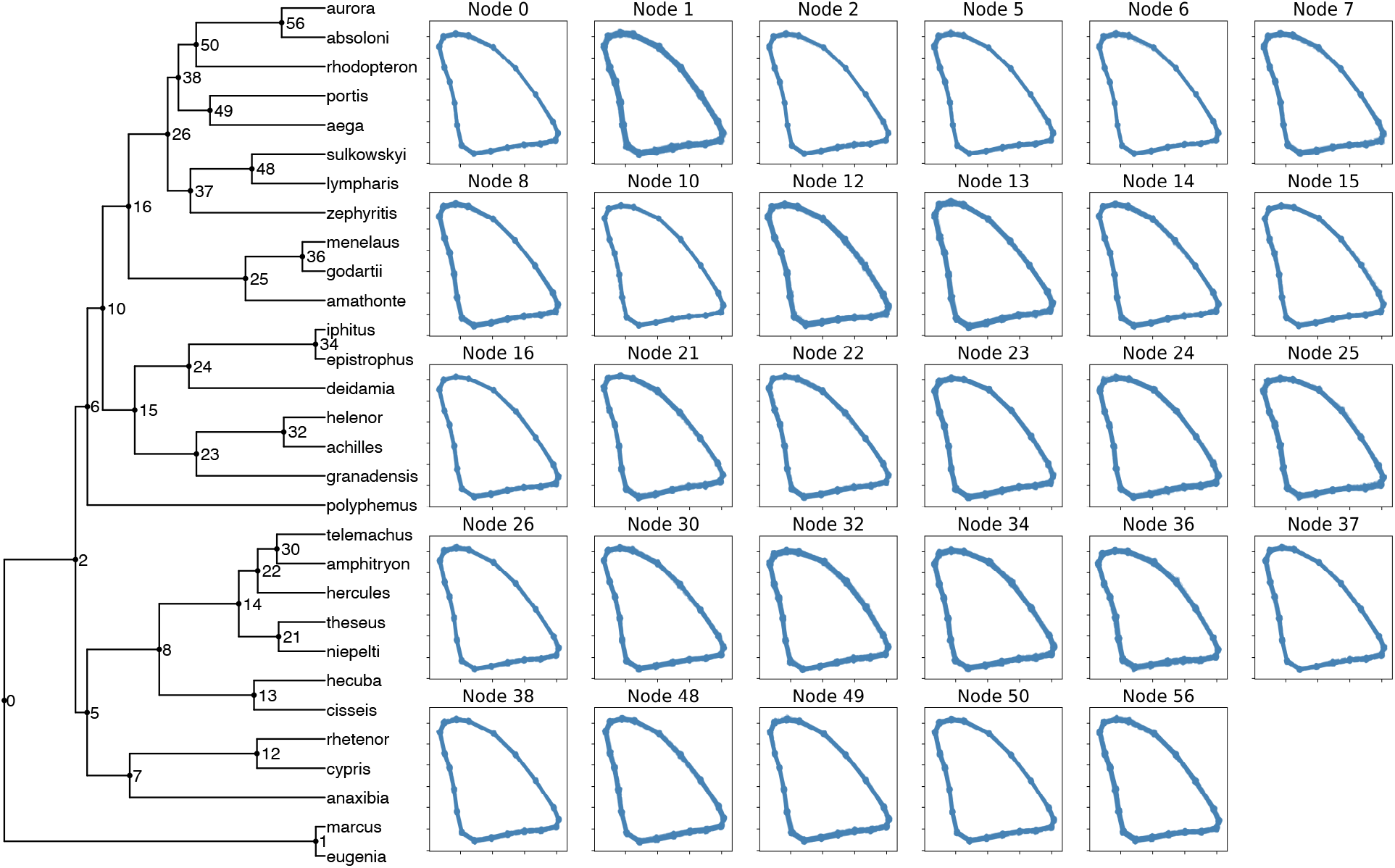
Figure showing the posterior distribution over ancestral states. On the left we show the phylogeny for the *Morpho* butterflies obtained form Chazot et al. (2021) with labelled internal nodes. On the right we show the posterior obtained from each internal node in the phylogeny. Each plot is labelled with a number corresponding to the numbers indicated in the phylogeny.

Lastly, we visualize posterior samples of the kernel parameters *α* and *σ* in Figure 12. As can be seen from the trace plot and 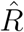 value the MCMC chains seem to have converged. For *α* parameter we estimate the posterior mean to be 0.0026 with 95% credible interval (0.002, 0.0032). Similarly, for *σ* we estimate the posterior mean to be 0.3434 with 95% credible interval (0.2648, 0.4292).

**Figure 12.**
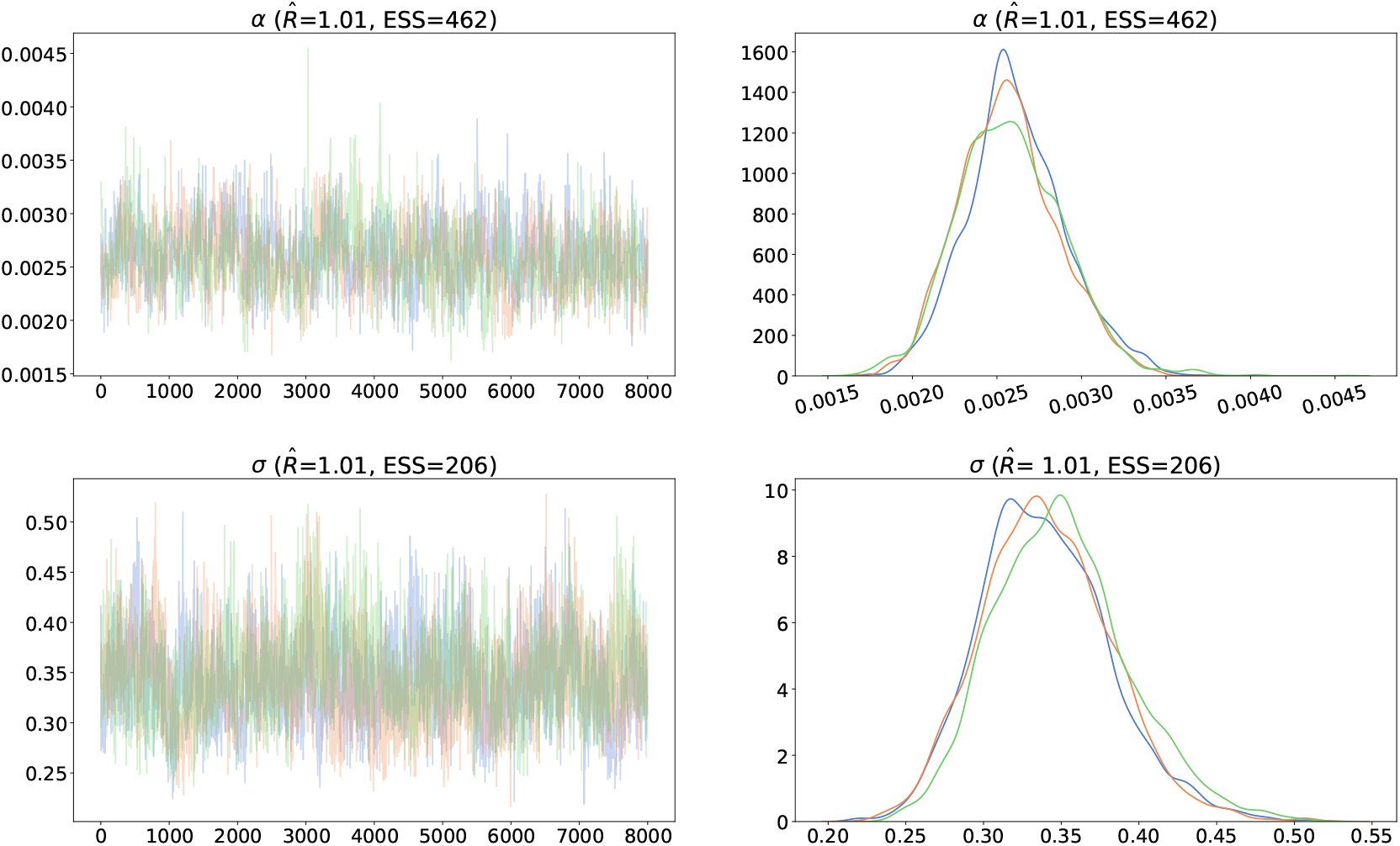
Figure showing the posterior distribution obtained for the different kernel parameters *α* and *σ*. Colors indicate the result for 3 independent chains. Leftmost column shows trace plots and rightmost column shows the marginal posterior densities for the different MCMC chains. Trace plots and density plots are shown after removal of burnin (first 2000 iterations). 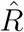 refers to an improved version of the Gelman-Rubin convergence diagnostics (Vehtari et al., 2021). ESS is the effective sample size.

## Discussion

### A Parametric Model of Shape

In this manuscript, we develop a framework for parametric stochastic shape model inference in phylogenetics, where the spatial autocorrelation among landmarks is built into the model. The parametric model we apply imposes a certain structure on the evolution of the landmarks, which has a number of distinct advantages. Firstly, it ensures that the shapes at all times are true shapes that do not self-intersect. In addition, it circumvents the large *p* small *n* problem. It does so because the covariance matrix of the model only depends on the current position of the landmarks and two parameters; *α* and *σ*. This is the case no matter how many landmarks we consider. Another advantage is that, when analyzing biological shapes, the parameters we infer are biologically meaningful. That is, we may interpret *α* as the evolutionary rate parameter and *σ* as a measure of biological integration (i.e. a measure of how correlated the landmarks in a shape are). When applying our model all relevant information about the process of shape evolution is contained in these two parameters. A challenge when applying the parametric shape model is that the diffusion term in the stochastic model becomes state-dependent i.e. depends on the current position of the landmarks. This is what makes inference harder and why we need the BFFG framework in order to infer the posterior distribution. Lastly, we note that our method makes the strong assumption of an isotropic Gaussian kernel on the covariance structure. In real biological data, this assumption may not be satisfied as certain parts of the morphology might evolve faster than others, or have varying correlation structure. The model we derive here might be viewed as a basic null-model for analyzing shape, that may form the foundation for testing hypothesis regarding covariance structures in future studies. We have not pursued such extensions to the method here, but focused on demonstrating the feasibility of incorporating explicit shape diffusion into the phylogenetic analysis of morphology

### Computational Considerations

A limitation of the applicability of the shape aware model is the computational complexity of the inference procedure. To give a concrete example, the analysis of the butterfly data set consisting of 30 species and 20 landmarks in 2D described in subsection *“Analysis of Morpho Butterflies”* took approximately 5 days and 5 hours to run on an Intel workstation using a single core. This reflects the current implementation, the computation will be able to run in significantly less time in future implementations. In the current implementation we propose first a path, then the *σ* parameter, and finally the *α* parameter for each iteration of the Metropolis-Hastings algorihtm. Each proposal is based on a backward filtering step followed by a forward guiding step. The most computationally heavy part of each proposal is the backward filtering step as this step requires inverting a *nd × nd* matrix *ζ/δ* times with *n* being the number of landmarks, *d* the dimension of the landmarks, *δ* the stepsize and *ζ* the total branch length in the tree. That is, the running time for the currently implemented MCMC algorithm is *O*(*ζ/δ* · (*n* · *d*)^3^). Although we have tried to make the current implementation fast, we still believe that there is much more work to be done in order to increase the computational speed of the inference procedure. In particular, we note that the indicated computational cost provides a worst case scenario in which no numerical linear algebra optimizations have been applied yet. In subsequent work we will explore such optimizations, which indicate that both the computational cost and storage requirements can be significantly reduced. In addition, we believe more can be done in order to properly benefit from the parallelization and Just-In-Time compilation available in Jax (a Python library). Alternatively, the code could also be implemented in C++ which would also be expected to lead to an important increase in speed.

### Algorithmic Improvements

In this paper, we infer one set of parameters *α* and *σ* on the entire tree. A natural extension of the MCMC algorithm would be to implement a version of the MCMC algorithm allowing for estimating different parameters on each branch in the phylogenetic tree. Increasing the number of parameters would require an increase in the number of backward passes in the MCMC algorithm. As backward passes are the most computationally heavy step in the algorithm, we would need a significantly faster implementation. In this study, we have found that the MCMC algorithm converges very slowly for specific parameter settings, particularly when considering the full phylogeny from Chazot et al. (2021). A very likely reason for this is that the current MCMC implementation is based on global updates, i.e., the entire stochastic path is updated at the same time. This is the case as the current implementation of the BFFG framework only allows for global updates. Therefore, an important topic for future research would also be developing a MCMC algorithm which could be based on local updates as this will very likely reduce the convergence problems and pave the way for applying this kind of model to analyze large phylogenies and estimate many parameters.

In this study we pursue a very simple strategy when it comes to inferring the posterior distribution over the root. We simply add an edge, connecting the root to a super root, and use a pre-selected shape as the value in the super root. This is done due to the difficulty of defining prior distributions for shapes. The shape process does not generate an easily defined stationary distribution of shapes. In the analyses conducted in this paper, we used the phylogenetic mean of the data as the shape in the super root. Furthermore, in the current implementation we do not distinguish between the parameters on the root to super root edge and the parameters governing the process in the rest of the tree. An essential improvement of the algorithm would therefore be to distinguish between the process on these two sets of edges to reduce the possible biases in the inferences of parameters stemming from the use of a super root.

### Future Directions

In this paper we have applied recent advances in stochastic shape analysis to model the evolution of biological shapes. To our knowledge, this paper is the first to apply shape-aware stochastic models to do evolutionary modeling of morphological data. As the shape-aware stochastic models allow us to do stochastic character mapping on the shapes themselves by directly modeling the trajectories of each landmark we believe that this type of model represents a promising modeling framework when it comes to analyzing biological shapes in an evolutionary context. In addition, the framework underlying this kind of model is general enough to also include full shapes, i.e., curves. Therefore, the work presented in this paper may also be considered a step towards transitioning from the analysis of landmark shapes to the analysis of full shapes in the context of evolutionary biology.

An essential part of analyzing empirical shape data is aligning the shapes i.e. positioning the shapes in a shared coordinate system. This is typically done using Procrustes analysis. Procrustes alignment minimizes the distances between shapes in terms of rotation, translation, and scaling and, therefore, will likely lead inference methods to underestimate the true rate of evolution. In fact, in the context of our stochastic model we have found that Procrustes aligning the data seems to result in both parameters getting slightly underestimated compared to estimates based on non-aligned data (see Supplementary Material Fig. S32 and S33). Although our stochastic shape model arguably is a more realistic model of shape evolution than previous methods, it still models translation, rotation and scaling, and does not take into account that shapes have been aligned. Not accounting for the effect of Procrustes alignment is a drawback of our method that is shared by all current stochastic methods. An important topic for future research is the development of models which allow for joint alignment and evolutionary analysis of shapes.

In this paper numerical discretisation was done using Euler discretisation, which is the very simplest method one could use. Specifically, we have used a stepsize of 0.05 for both simulating data sets and estimating the posterior distribution. However, as Euler discretisation is not unconditionally stable we acknowledge that for empirical data sets we are not guaranteed that a stepsize of 0.05 will be sufficiently small to properly approximate the shape process. Therefore, depending on the data set, more elaborate discretisation schemes might be preferable. To date, the authors are not aware of any quantitative results comparing such schemes for the model we consider here.

The stochastic model we apply in this study assumes a constant correlation function at every point in space and time. It could perhaps be considered a null model against which future, more complicated models, could be tested. Extending the model to directly incorporate biological hypotheses is an important topic for future research. For instance, the model could be extended to model specific subregions of a shape under selection, i.e., where, during certain periods of time, the evolutionary dynamics follows other parameters than the rest of the shape. We also imagine extending the model such that it allows for identification of evolutionary modules in the shape i.e. different subregions where all landmarks seem to be more correlated than what we would expect only based on the distances between the landmarks.

Another exciting avenue for future research includes combining our framework with stochastic models of other traits such as DNA or environmental variables. Such joint models would help identify correlations between the rate of morphological evolution and other traits, such as DNA substitution rates. As modern evolutionary biology is based on analysis of large phylogenies, the greatest obstacle, when it comes to developing such joint models, is the computational speed of our inference procedure. Computational challenges similarly limit the method, at the moment, from becoming a vehicle for estimation of phylogenetic topology.

## Conclusion

This study introduces a new framework for evolutionary analysis of shape data that is based on an explicit model of shape evolution. We use it here to illustrate how simple parameters representing evolutionary rate and morphological integration can be estimated, and to obtain credible sets of ancestral reconstructions of shape. However, the advantage of this framework goes further than that. It allows for explorations of evolutionary hypotheses regarding the processes governing evolutionary shape deformations in an explicit probabilistic framework. Studies of DNA evolution have long benefited from such explicit models for testing hypotheses regarding evolution of DNA. Shape explicit models will allow similar explorations of the laws and processes governing shape evolution.

## Funding Acknowledgments

This work was supported by a research grant (VIL40582) from VILLUM FONDEN and the Novo Nordisk Foundation grants NNF18OC0052000, NNF24OC0093490.

## Acknowledgments

Firstly, we wish to thank Anders Munch, Dean Adams and Kasper Munch for thoroughly reading our manuscript and sharing their thoughts and ideas. We also thank Elizabeth Baker and Ricardo Ely for fruitful and ongoing discussions. Finally, we thank Nicolas Chazot for making his data publicly available, Michael Severinsen for removing the background from the butterfly images from GBIF and the developers behind ETE 3.0 (Huerta-Cepas et al., 2016), ArviZ (Kumar et al., 2019), and Jax (Bradbury et al., 2018) for developing Python packages that have been essential for the development of our code.

## Supplementary Material

All scripts necessary to reproduce the analyses reported in this study can be accessed through the Zenodo link: https://doi.org/10.5281/zenodo.15109856. In addition, the code is also available on GitHub: https://github.com/sofiastroustrup/SPMS/tree/main.

