## Appendix for "Stochastic Phylogenetic Models of Shape"

### Appendix 1. Infinite-dimensional Formulation of the Shape SDE

As described, the shape SDE in equation (2) is related to *Kunita flows*. In section *"Finite-dimensional Formulation of the Shape SDE"* we looked at solutions in a point-wise fashion, which is natural from the point of view of numerical discretization. In this section we briefly describe how a Kunita flow can be considered more abstractly as an SDE on a function space, in particular diffeomorphisms. This formulation shows that the process exists independently of the chosen grid, and it opens up for alternative implementations.

The more general formulation of our shape Kunita flow given in equation (3) is as follows. Let $Q$ be a Hilbert-Schmidt operator on the Hilbert space $H = L^2(\mathbb{R}^d, \mathbb{R}^d)$ (square-integrable maps from $\mathbb{R}^d$ to $\mathbb{R}^d$). It is then on integral form so there exists a kernel $k$ and $Q(f)(x) = \int_{\mathbb{R}^d} k(x,y)f(y)dy$. For $h \in H$, we furthermore define $(Qh)f(x) = \int_{\mathbb{R}^d} k(h(x) + x, y)f(y)dy$ and consider the $H$-valued SDE

$$X(t) = \int_0^t Q(X(t))dW(t), \qquad X(0) = 0$$

where $W(t)$ is a cylindrical Brownian motion. It can be shown that for solutions $X(t)$, the maps $y \mapsto X(t)(y) + y$ are diffeomorphisms for every time $t$. Such flows of diffeomorphisms are denoted Kunita flows (see, e.g., Kunita (1990) and Baker et al. (2024)). From this flow, we can recover the path of a particular landmark or point $x$ as $t \to X(t)(x)$. Due to $Q$ being a convolution operator, the SDE convolves the diffeomorphism $X(t)$ with the background noise $W(t)$, which amounts to a random perturbation of the diffeomorphism.

### Appendix 2. Reflected Gaussian Proposal

We note that the algorithm for sampling from a reflected Gaussian described below is defined for one dimensional variables.

**Algorithm 1** Sampling from a Reflected Gaussian

```
κ° ~ N(κ, τ²)
while κ° < a or κ° > b: do
    if κ° < a then
        κ° := 2a − κ°
    else
        if κ° > b then
            κ° := 2b − κ°
        end if
    end if
end while
```

### Appendix 3. Simulation-Based Calibration

For a Bayesian model the data averaged posterior reduces to the prior distribution when the data are simulated from the Bayesian model i.e.

$$\pi(\theta) = \int \pi(\theta|\tilde{y})\pi(\tilde{y}|\tilde{\theta})\pi(\tilde{\theta})d\tilde{y}d\tilde{\theta}$$

where the parameter $\tilde{\theta}$ is sampled from the prior distribution over the parameter i.e. $\tilde{\theta} \sim \pi(\theta)$ and the data is simulated from the generative model i.e. $\tilde{y} \sim \pi(y|\tilde{\theta})$. Note that $\pi(y|\tilde{\theta})$ is the distribution of the data $y$ given a certain parameter $\tilde{\theta}$ and $\pi(\theta)$ is the prior distribution over the parameter $\theta$. In our setting, $\tilde{y}$ is the observed shapes in the leaves of the phylogenetic tree, and $\tilde{\theta}$ denote the kernel parameters and the latent ancestral shapes within the tree. The fact that the data averaged posterior reduces to the prior, is the foundation for Simulaton-Based Calibration (SBC) (Talts et al., 2018) which we use to evaluate the computational aspects of our Metropolis-Hastings algorithm. We follow the approach described in Talts et al. (2018) and compare the data averaged posterior with the prior distribution using rank statistics. As shown in Talts et al. (2018) the rank statistics will be uniformly distributed if the data averaged posterior and the prior distribution are identical hence the Bayesian analysis has been correctly implemented. In (Talts et al., 2018) the rank statistic $r$, of the prior sample relative to the posterior sample is computed for a one-dimensional random variable, $f : \Theta \to \mathbb{R}$, as

$$r(\{f(\theta_1), \cdots, f(\theta_L)\}, f(\tilde{\theta})) = \frac{1}{L} \sum_{l=1}^{L} \mathbb{I}[f(\theta_l) < f(\tilde{\theta})]$$

where $f(\theta_l)$ is a sample from the marginal posterior distribution and $f(\tilde{\theta})$ is the true marginal value obtained from simulation. As the rank statistics are computed based on one-dimensional random variables, we assess histograms of rank statistics for all dimensions of all landmarks in all nodes as well as for each of the parameters.
