## Supplementary Material for "Stochastic Phylogenetic Models of Shape"

### Supplementary Material
### Stochastic Phylogenetic Models of Shape

#### 1 Butterfly images

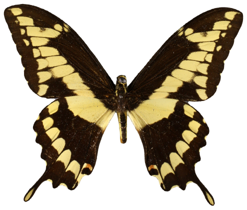
(a)

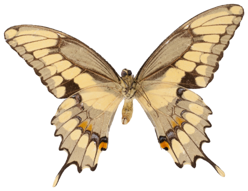
(b)

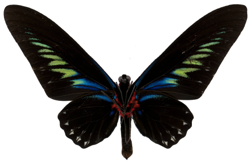
(c)

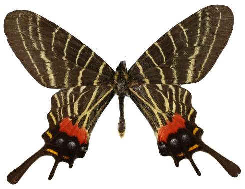
(d)

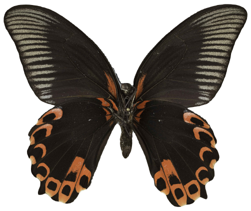
(e)

Figure 1

Subfigure **a**: GBIF ID = 1229919446, distributed under CC BY NC 4.0, Museum of Comparative Zoology, Harvard University, `http://mczbase.mcz.harvard.edu/guid/MCZ:Ent:135435`. We have removed the background from the image.

Subfigure **b**: GBIF ID = 1318834134, distributed under CC0 1.0, National Museum of Natural History, Smithsonian Institution. We have removed the background from the image.

Subfigure **c**: GBIF ID = 1322933267, distributed under CC0 1.0, National Museum of Natural History, Smithsonian Institution. We have removed the background from the image.

Subfigure **d**: GBIF ID = 1438640361, distributed under CC BY NC 4.0, Museum of Comparative Zoology, Harvard University, `https://mczbase.mcz.harvard.edu/guid/MCZ:Ent:135064`. We have removed the background from the image.

Subfigure **e**: GBIF ID = 1457664398, distributed under CC BY NC 4.0, Natural History Museum of Utah (UMNH), `https://ecdysis.org/collections/individual/index.php?occid=1399184`. We have removed the background from the image.

### 2 Supplementary Material for section *"MCMC Algorithm Yields Well Calibrated Credible Sets"*

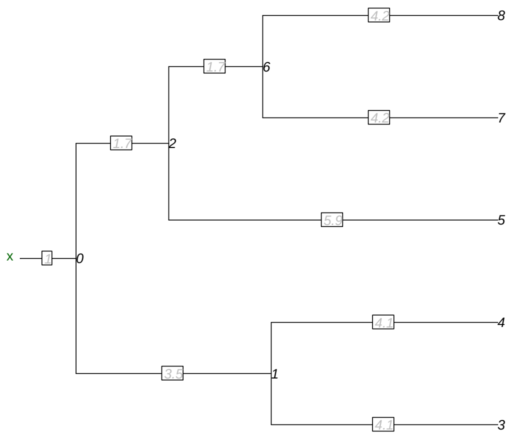


Figure 2: Figure showing the phylogeny used for simulation and inference in SBC. The branch lengths in the phylogeny are shown in grey on the specific branch and the root is denoted with $x$. This phylogeny is a subtree of the phylogeny from Chazot et al. (2021) where we have added a branch of length 1 going into the root. The number next to nodes in the phylogeny indicate the numbering used to identify specific nodes and correponds to the naming used in the Rhat plots and rank statistics plot.

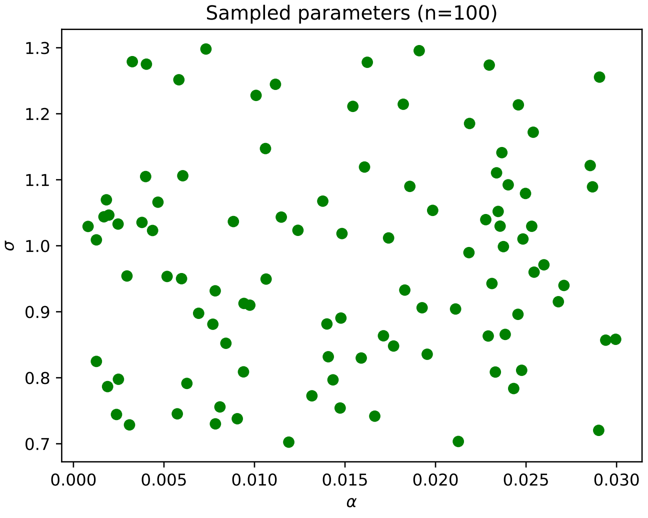


Figure 3: Paramters used for SBC.

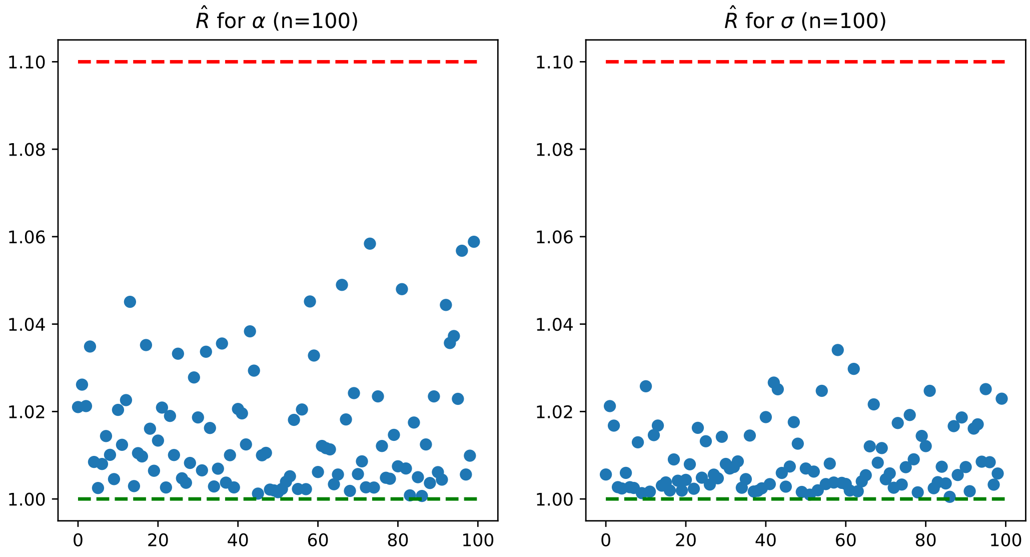


Figure 4: Figure showing the distribution of $\hat{R}$ for both kernel parameters. Index on x-axis corresponds to different data sets.

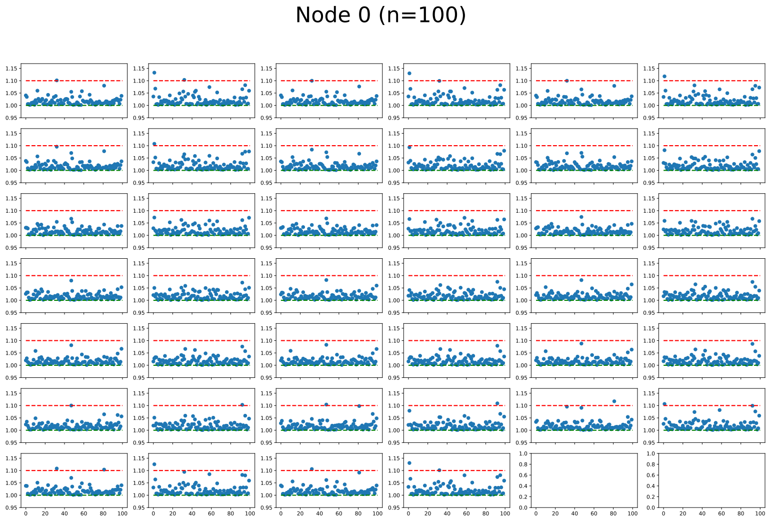


Figure 5: $\hat{R}$ for each dimension of node 0. Index on x-axis corresponds to different data sets. The node indices used are shown in supplementary Figure 2.

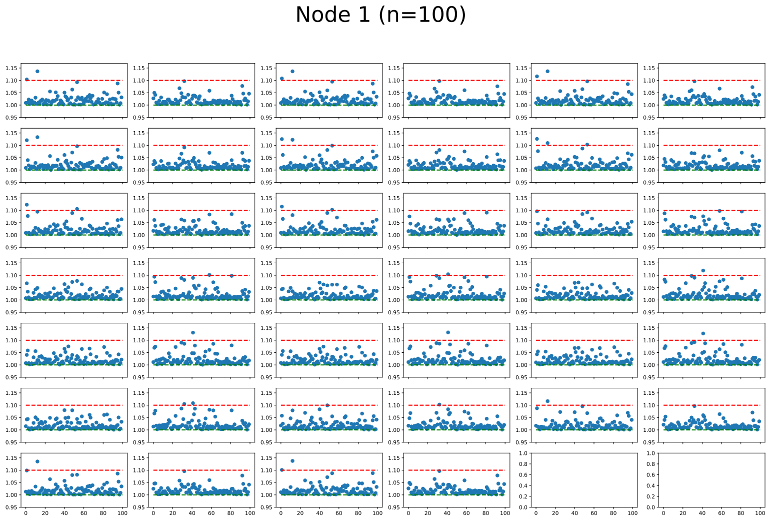


Figure 6: $\hat{R}$ for each dimension of node 1. Index on x-axis corresponds to different data sets. The node indices used are shown in supplementary Figure 2.

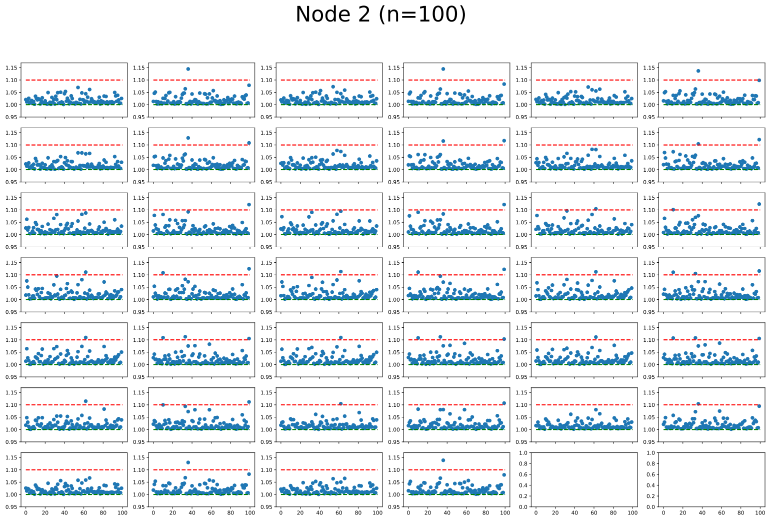


Figure 7: $\hat{R}$ for each dimension of node 2. Index on x-axis corresponds to different data sets. The node indices used are shown in supplementary Figure 2.

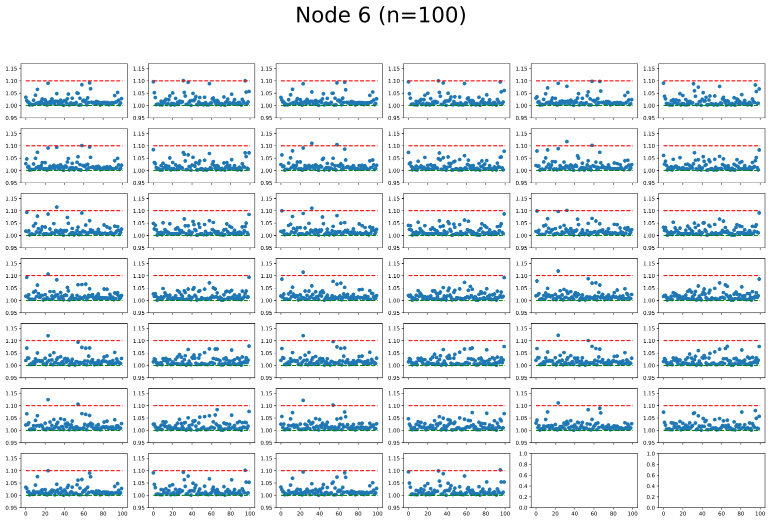


Figure 8: $\hat{R}$ for each dimension of node 6. Index on x-axis corresponds to different data sets. The node indices used are shown in supplementary Figure 2.

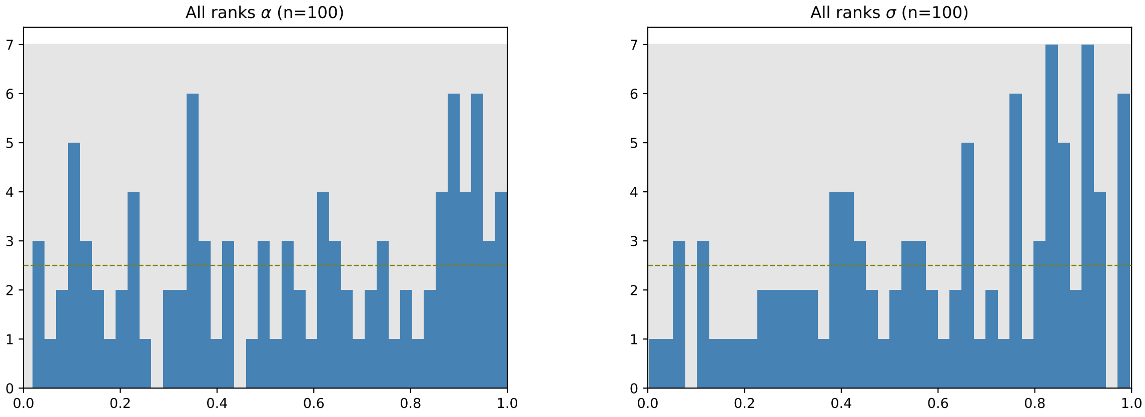


(a) (b)

Figure 9: Figure showing the distribution of ranks for the parameters of the shape process. Grey areas represent 99% of the variance we would expect under uniformity. Green dotted horizontal line represent the expected number of observations in each bin. **(a)** Rank distribution for parameter $\alpha$. **(b)** Rank distribution for parameter $\sigma$.

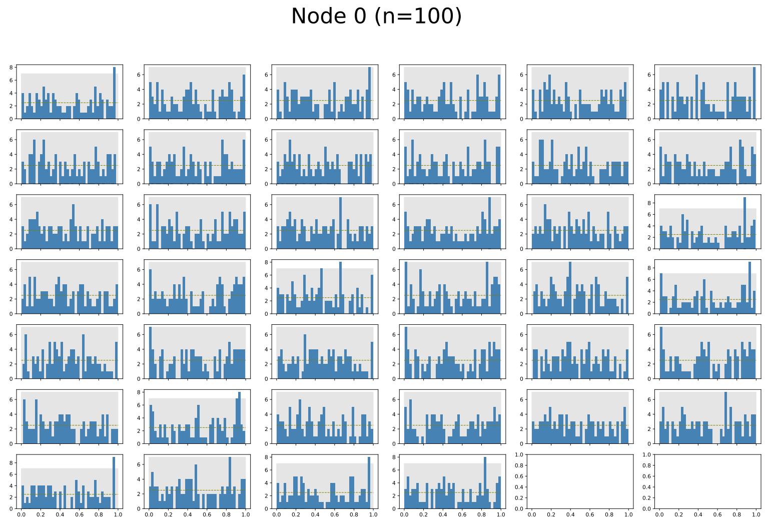


Figure 10: Distribution of rank statistics for all dimensions of node 0. The node indices used are shown in supplementary Figure 2.

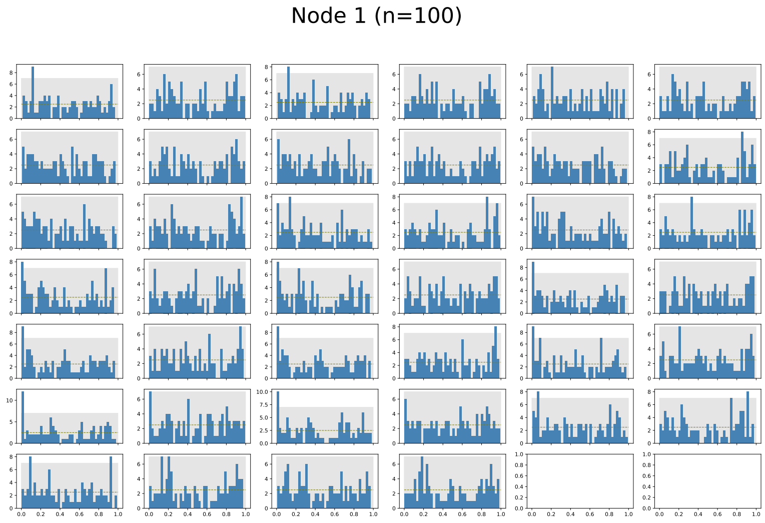


Figure 11: Distribution of rank statistics for all dimensions of node 1. The node indices used are shown in supplementary Figure 2.

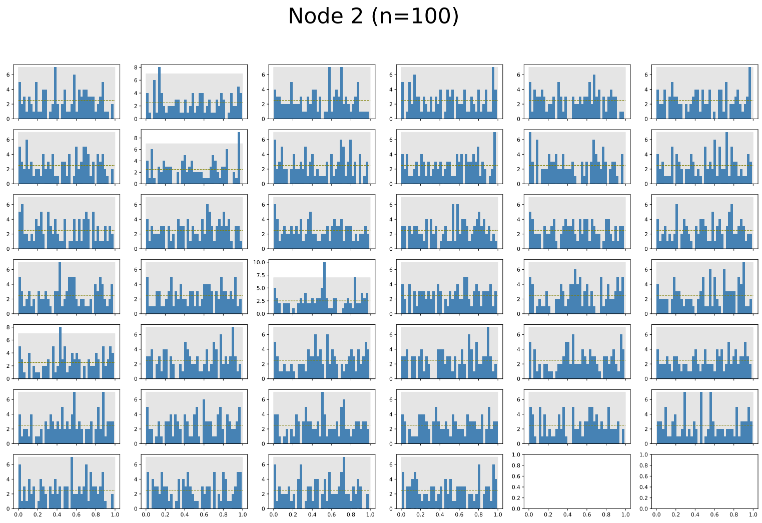


Figure 12: Distribution of rank statistics for all dimensions of node 2. The node indices used are shown in supplementary Figure 2.

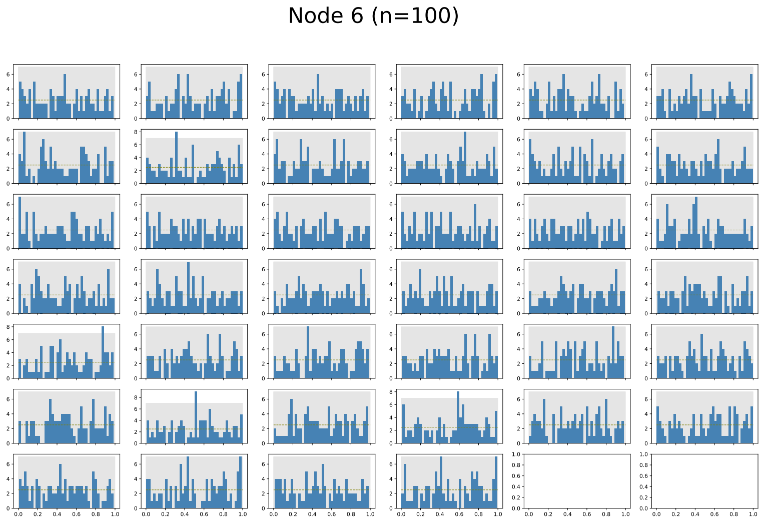


Figure 13: Distribution of rank statistics for all dimensions of node 6. The node indices used are shown in supplementary Figure 2.

### 3 Supplementary Material for section *"Root Estimation"*

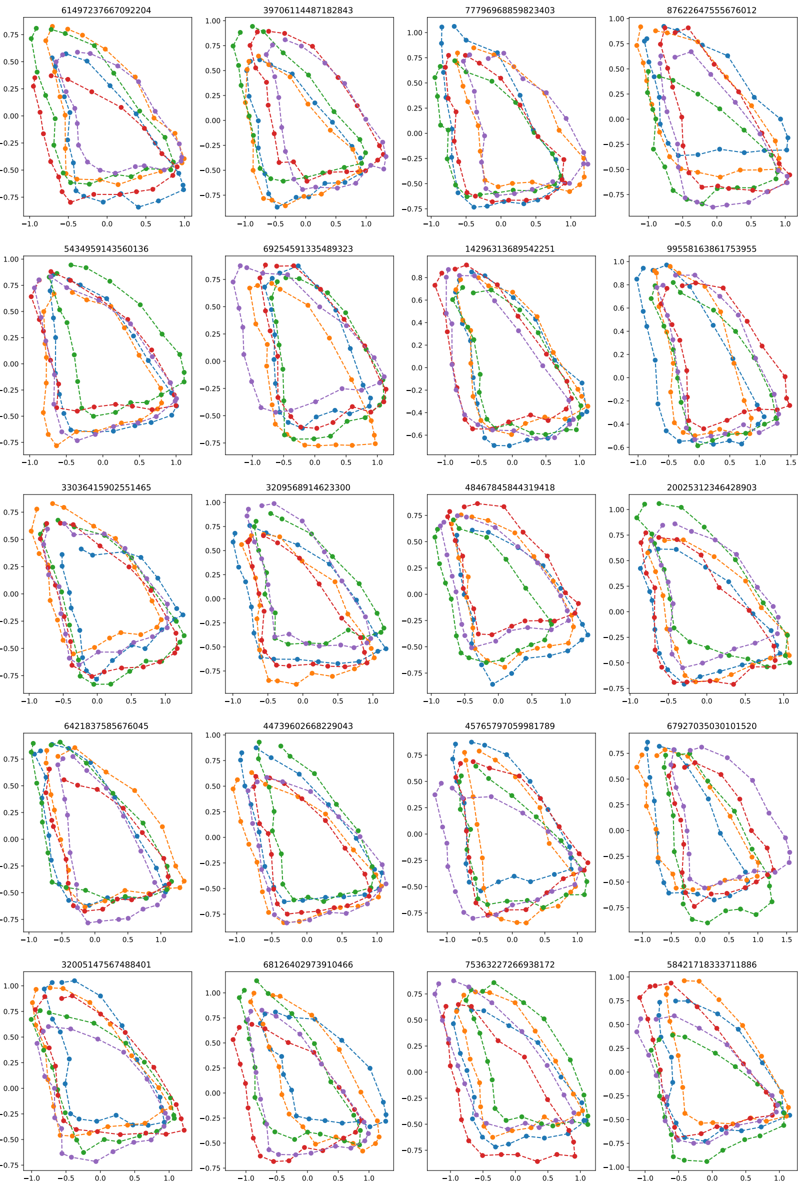


Figure 14: Visualization of the 20 simulated data sets used for evaluating root estimation. Data is simulated on a mixed topology with parameters $\alpha = 0.02$ and $\sigma = 0.9$. Colors refer to different tip states and title of each plot is the seed used for simulating the data.

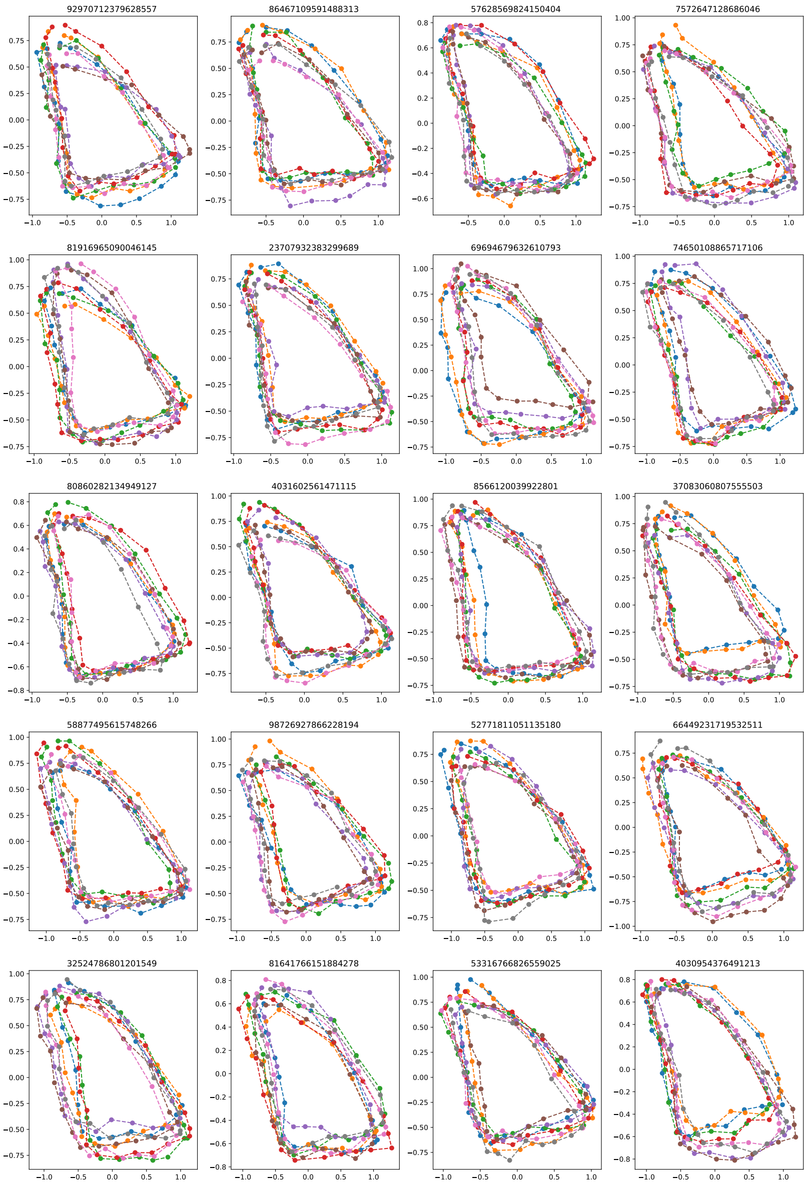


Figure 15: Visualization of the 20 simulated data sets used for evaluating root estimation. Data is simulated on the symmetric topology with parameters $\alpha = 0.025$ and $\sigma = 0.7$. Colors refer to different tip states and title of each plot is the seed used for simulating the data.

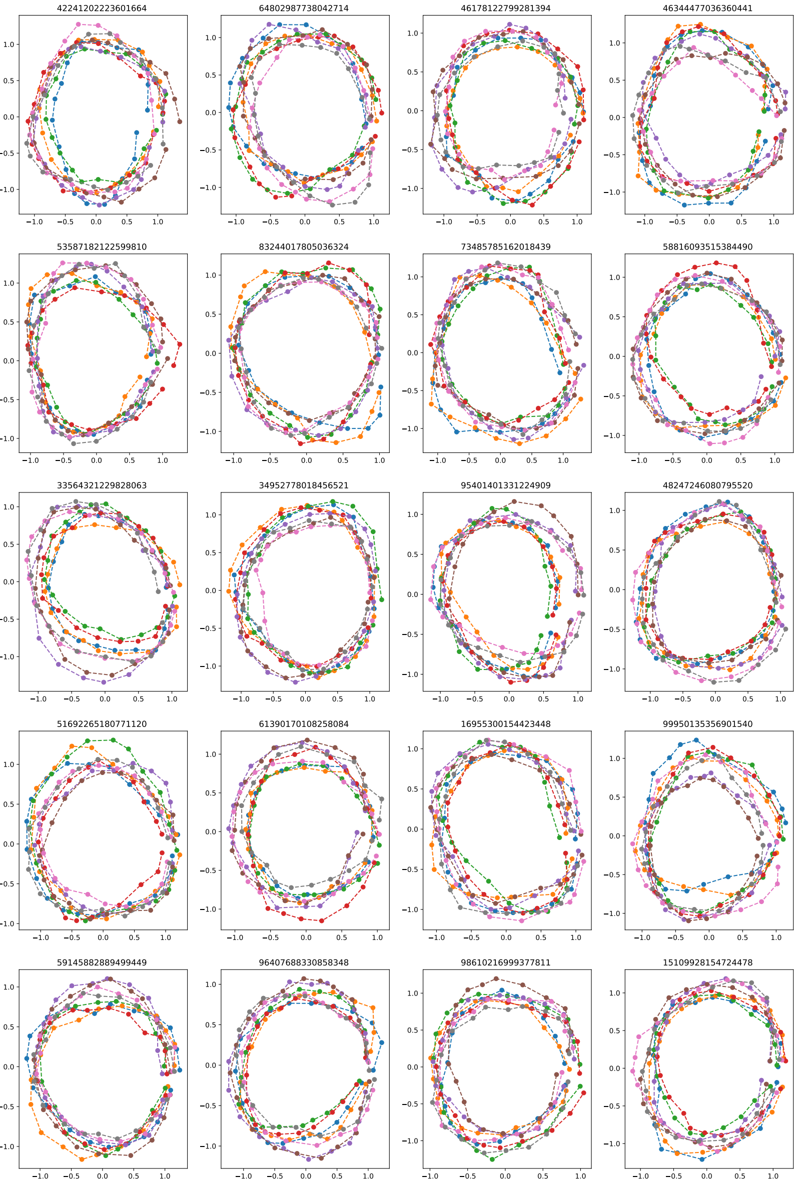


Figure 16: Visualization of the 20 simulated data sets used for evaluating root estimation. Data is simulated on the symmetric topology with parameters $\alpha = 0.04$ and $\sigma = 0.5$. Colors refer to different tip states and title of each plot is the seed used for simulating the data.

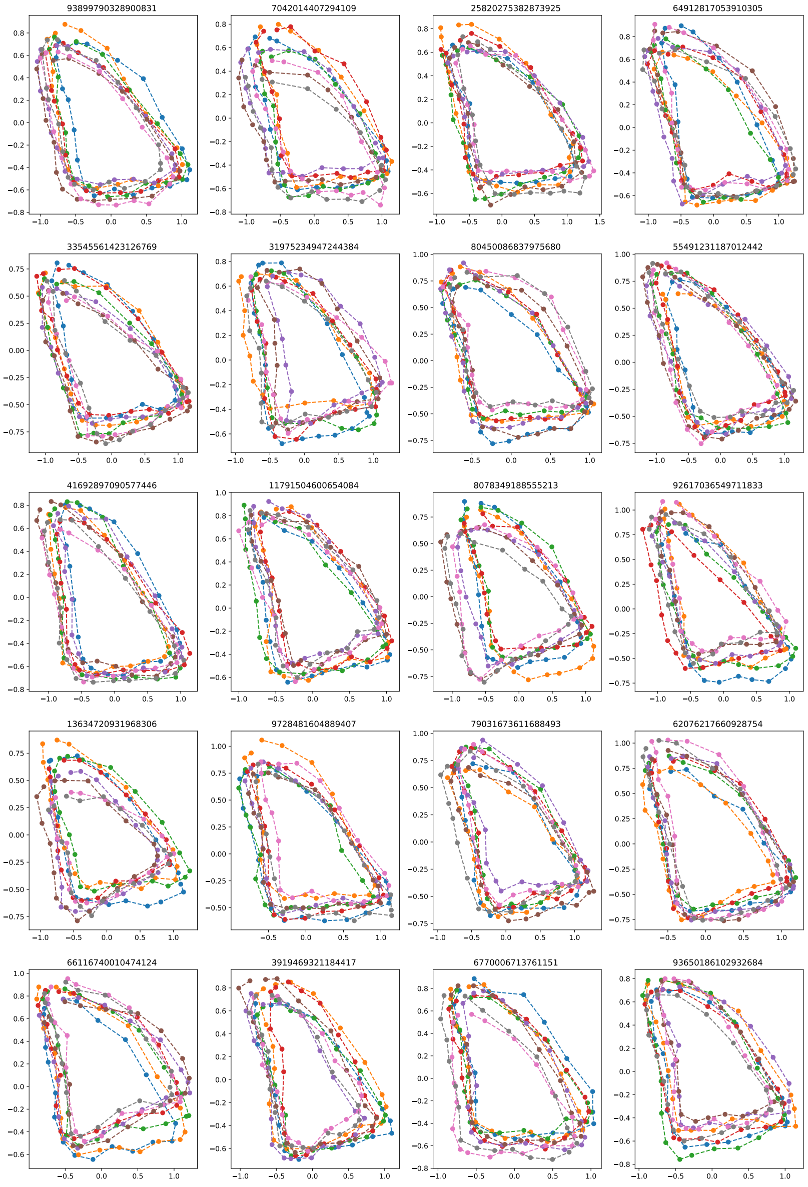


Figure 17: Visualization of the 20 simulated data sets used for evaluating root estimation. Data is simulated on the asymmetric topology with parameters $\alpha = 0.025$ and $\sigma = 0.7$. Colors refer to different tip states and title of each plot is the seed used for simulating the data.

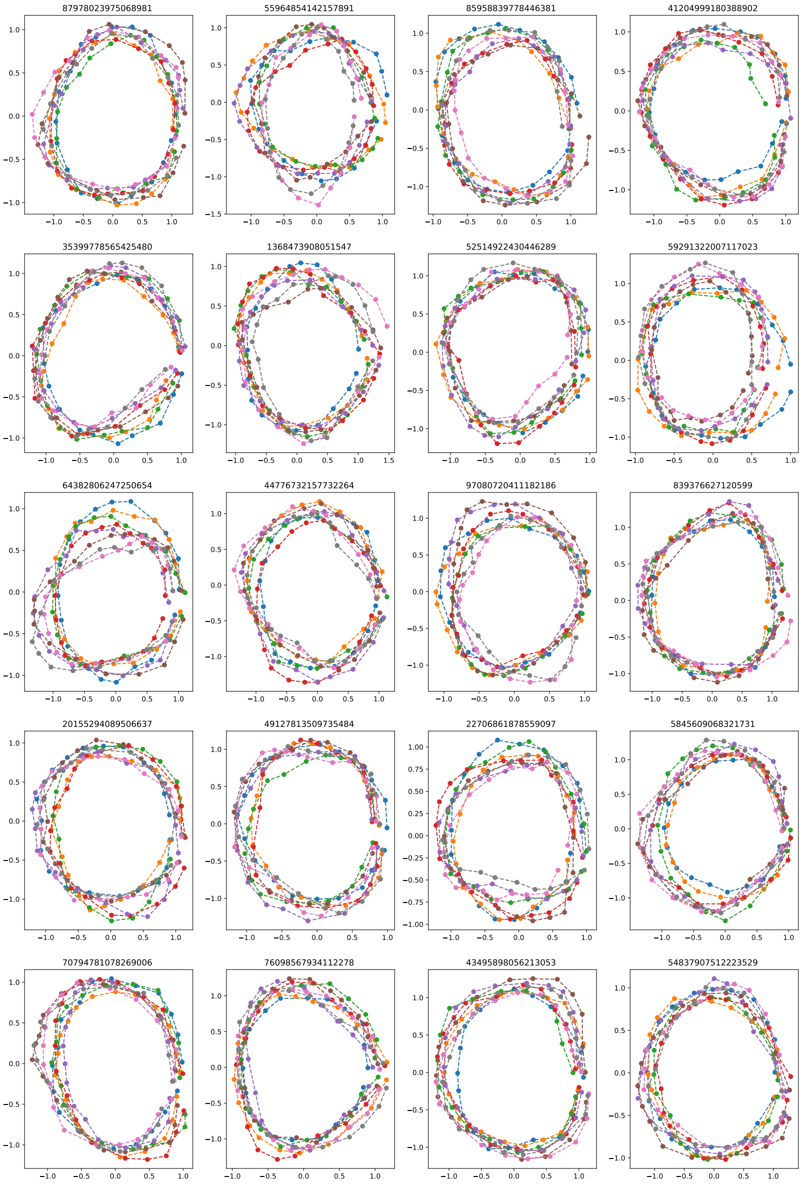


Figure 18: Visualization of the 20 simulated data sets used for evaluating root estimation. Data is simulated on the asymmetric topology with parameters $\alpha = 0.04$ and $\sigma = 0.5$. Colors refer to different tip states and title of each plot is the seed used for simulating the data.

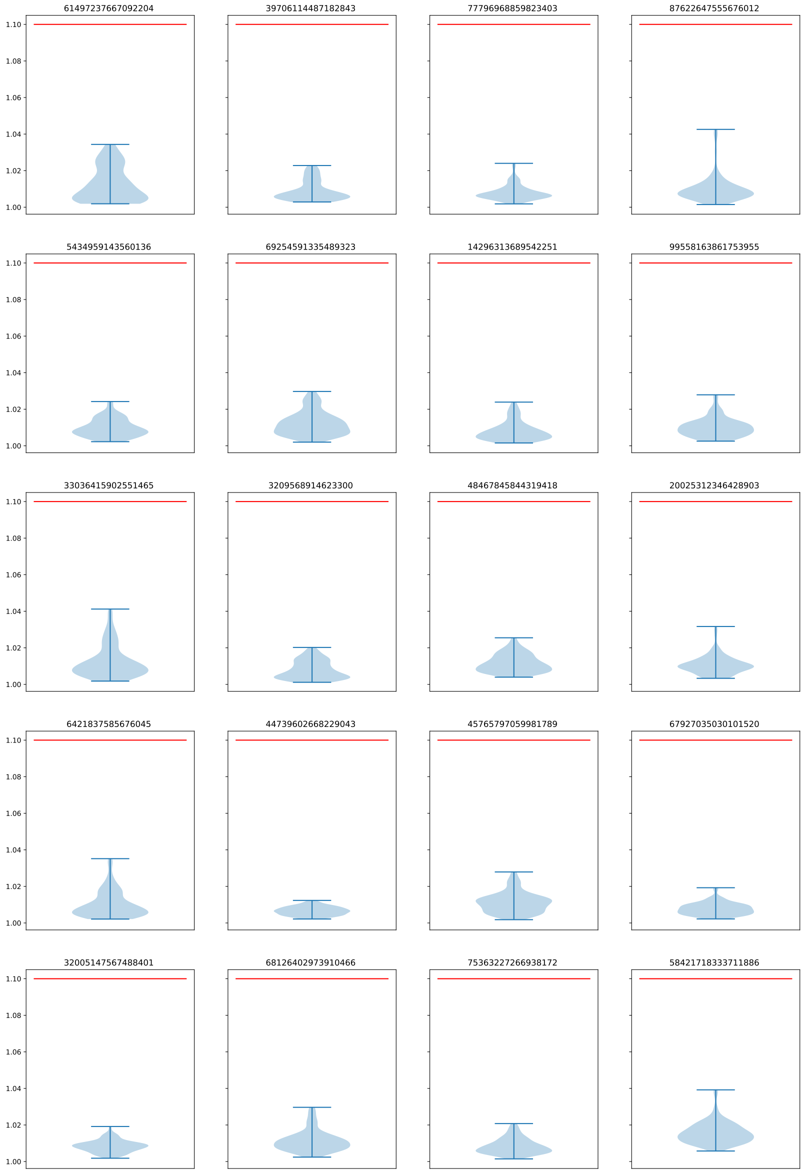


Figure 19: Visualization of convergence diagnostics for each of the 20 simulated data sets. Data sets where simulated on the mixed topology with the root shape being a butterfly forewing. Each plot show the distribution of 3200 $\hat{R}$ values describing all dimensions of all landmarks in all internal nodes and the parameters $\sigma$ and $\alpha$.

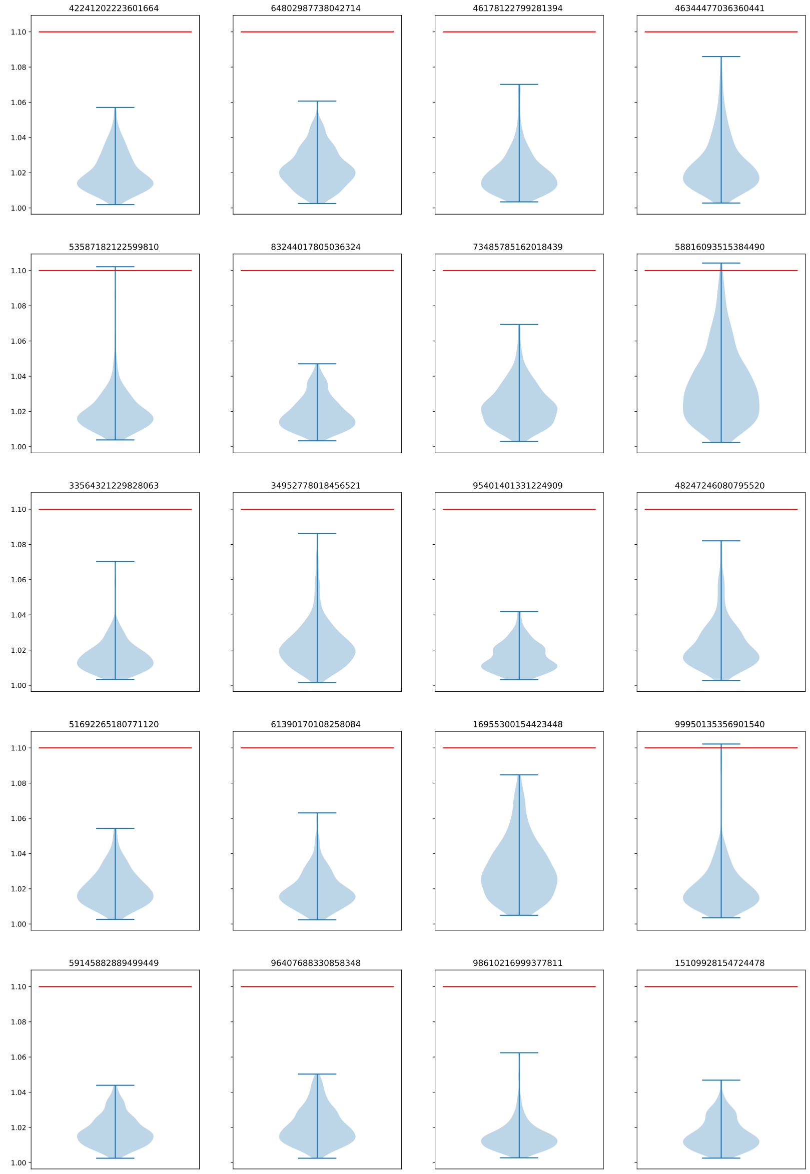


Figure 20: Visualization of convergence diagnostics for each of the 20 simulated data sets. Data sets where simulated on the symmetric topology with the root shape being a circle. Each plot show the distribution of 5600 $\hat{R}$ values describing all dimensions of all landmarks in all internal nodes and the parameters $\sigma$ and $\alpha$.

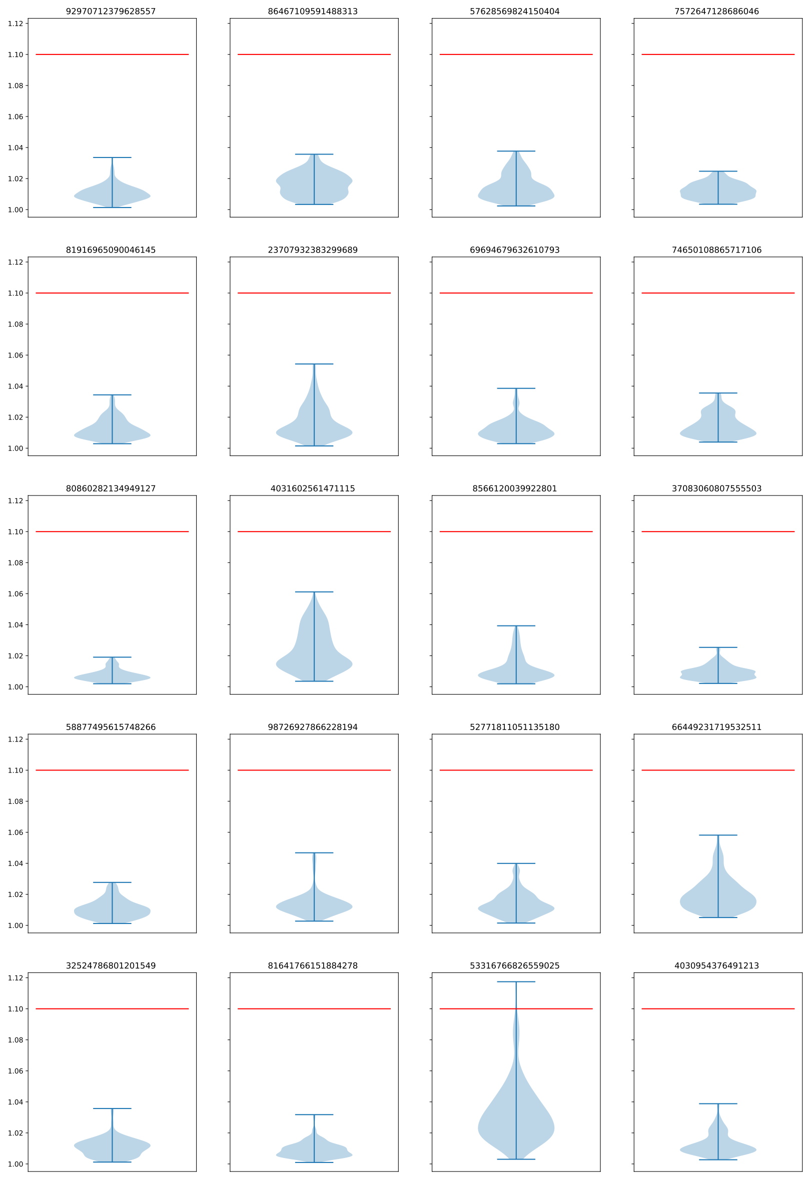


Figure 21: Visualization of convergence diagnostics for each of the 20 simulated data sets. Data sets where simulated on the symmetric topology with the root shape being a butterfly forewing. Each plot show the distribution of 5600 $\hat{R}$ values describing all dimensions of all landmarks in all internal nodes and the parameters $\sigma$ and $\alpha$.

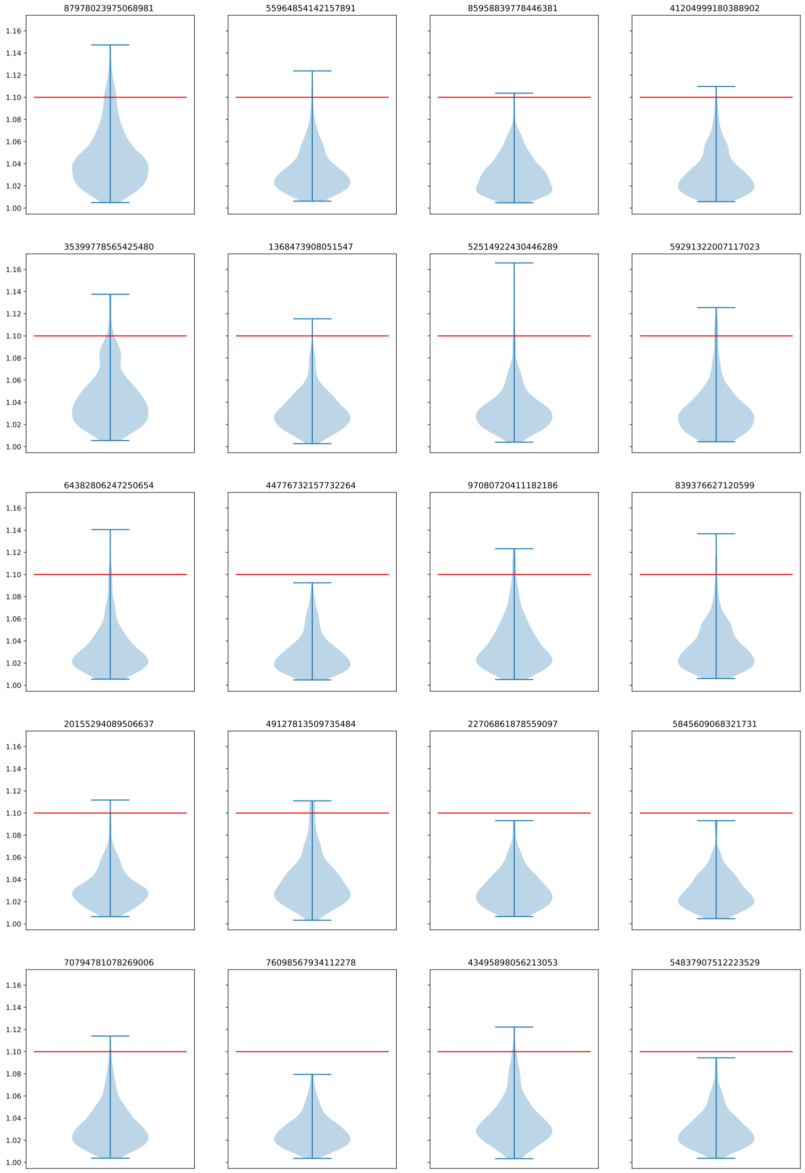


Figure 22: Visualization of convergence diagnostics for each of the 20 simulated data sets. Data sets where simulated on the asymmetric topology with the root shape being a circle. Each plot show the distribution of 5600 $\hat{R}$ values describing all dimensions of all landmarks in all internal nodes and the parameters $\sigma$ and $\alpha$.

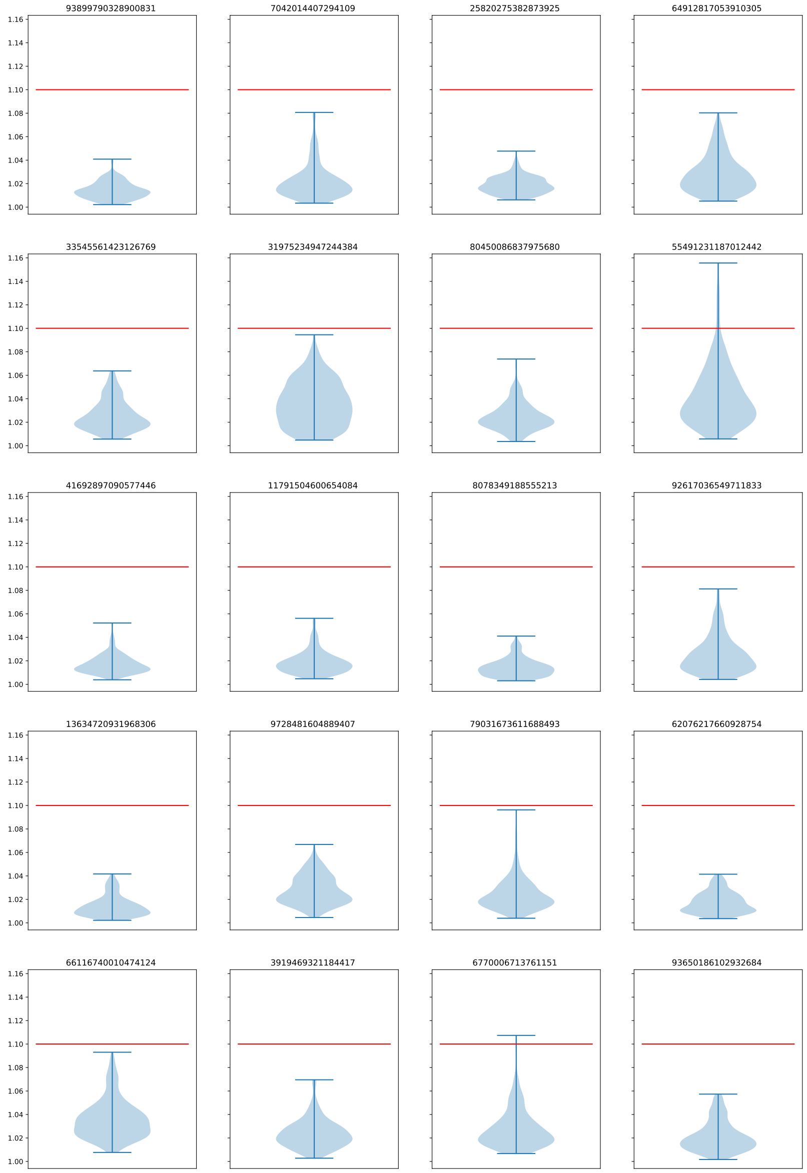


Figure 23: Visualization of convergence diagnostics for each of the 20 simulated data sets. Data sets where simulated on the asymmetric topology with the root shape being a butterfly forewing. Each plot show the distribution of 5600 $\hat{R}$ values describing all dimensions of all landmarks in all internal nodes and the parameters $\sigma$ and $\alpha$.

### 4 Supplementary Material for section *"Probabilistic Ancestral Reconstruction"*

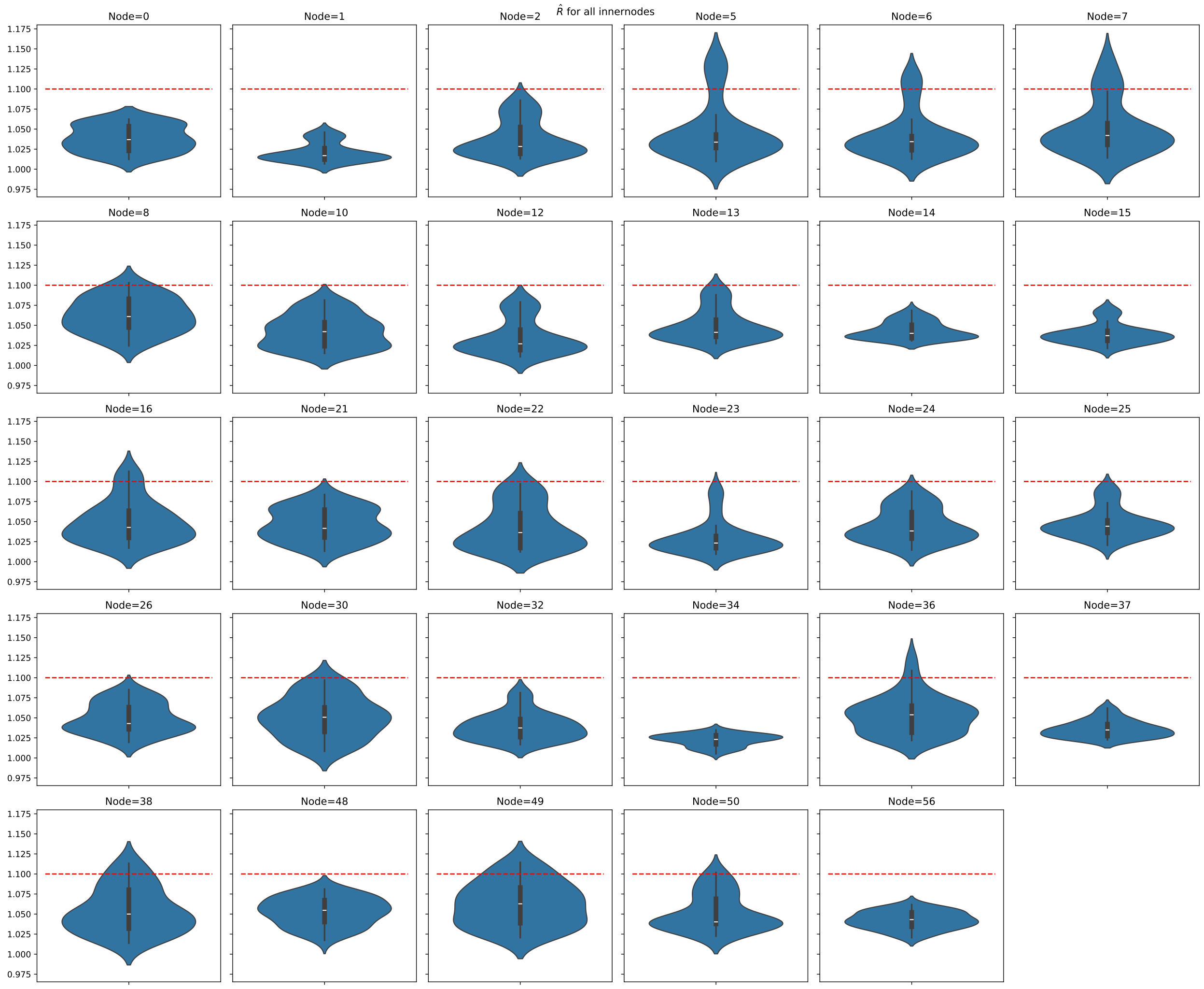


Figure 24: Distribution of $\hat{R}$ for all 40 dimension of each internal node.

### 5 Supplementary Material for section *"Analysis of Morpho Butterflies"*

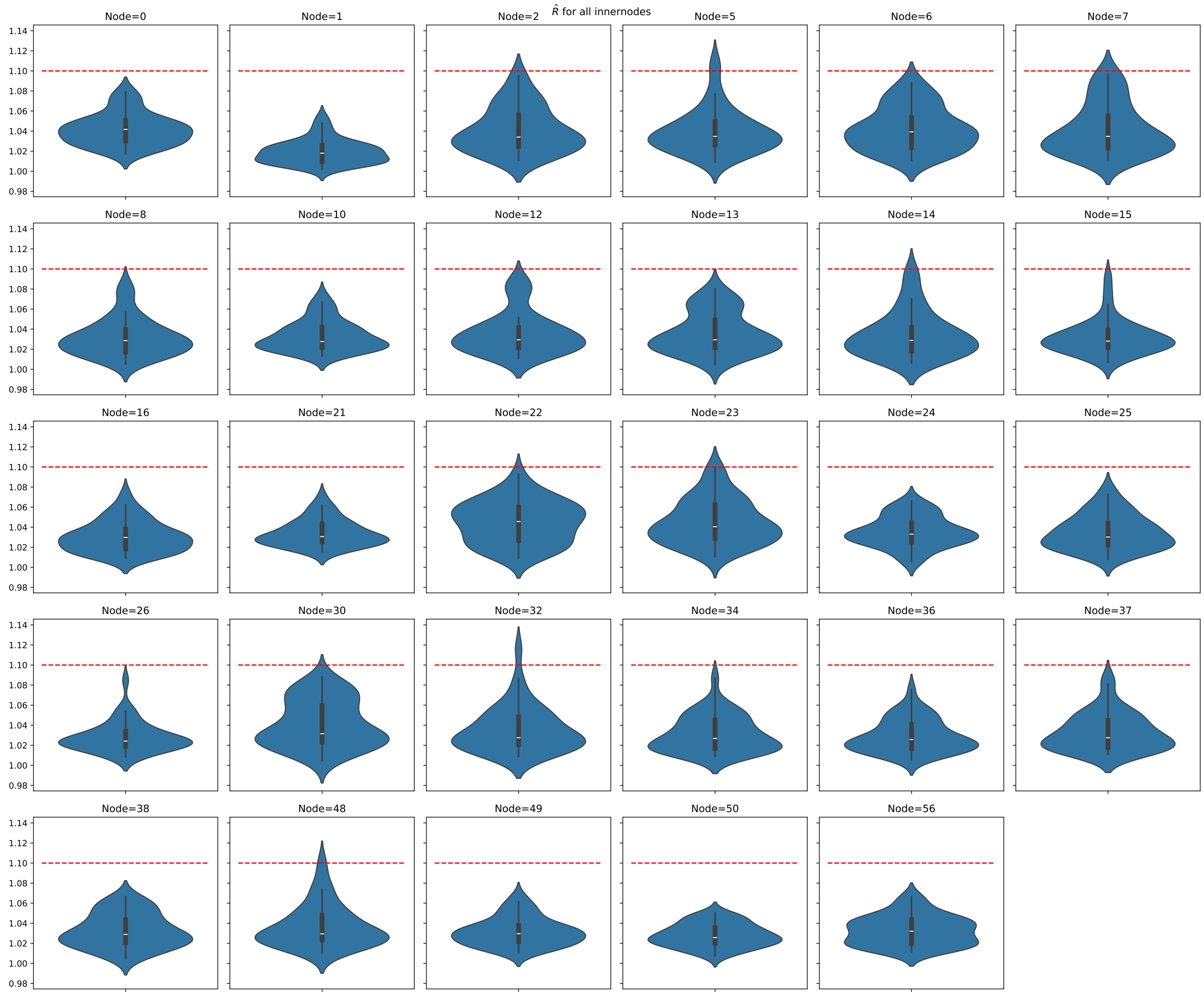


Figure 25: Distribution of $\hat{R}$ for all 40 dimension of each internal node.

### 6 Supplementary Material for section *"Shape-Aware Model Yield Meaningful Uncertainty Estimates"*

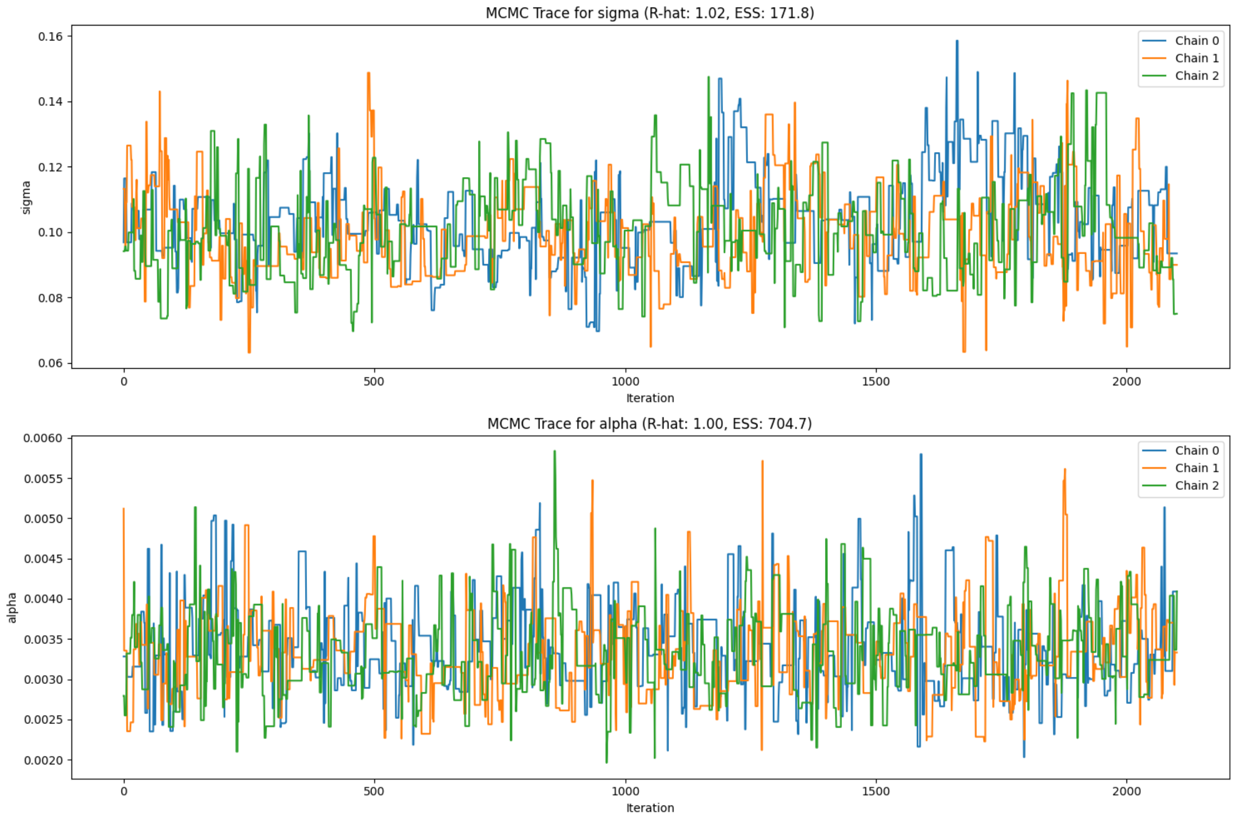


Figure 26: Trace plots for estimation of the kernel parameters, $\alpha$ and $\sigma$. Colors indicate the result of three independent chains. Trace plots are shown after removing a burnin of 30%. $\hat{R}$ refers to an improved version of the Gelman-Rubin convergence diagnostics. ESS is the effective sample size.

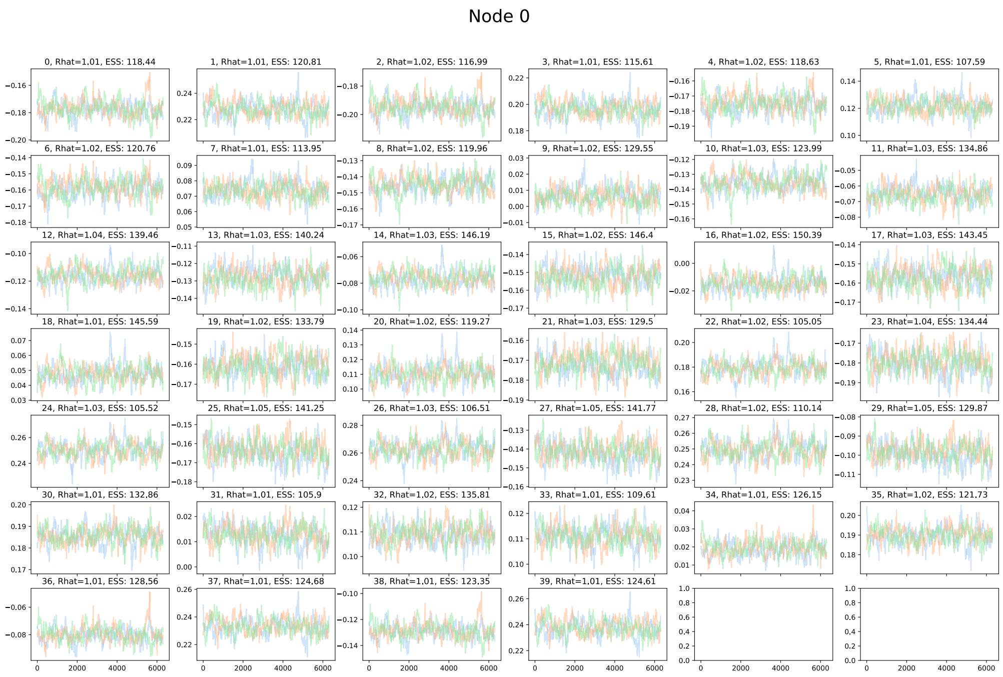


Figure 27: Trace plots for estimation of ancestral states. Colors indicate the result of three independent chains. Trace plots are shown after removing a burnin of 30%. $\hat{R}$ refers to an improved version of the Gelman-Rubin convergence diagnostics. ESS is the effective sample size.

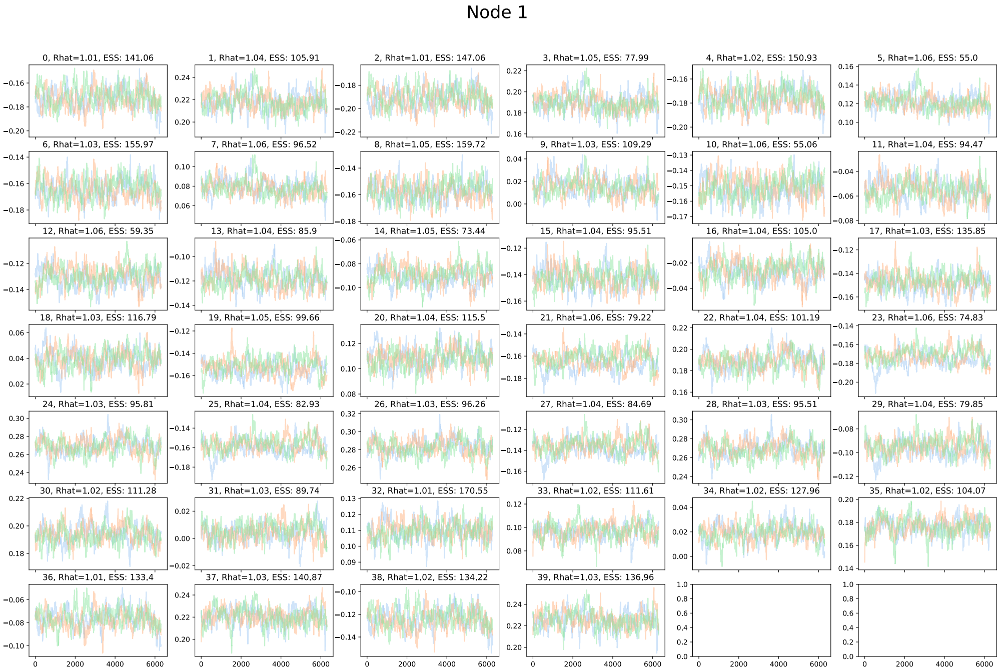


Figure 28: Trace plots for estimation of ancestral states. Colors indicate the result of three independent chains. Trace plots are shown after removing a burnin of 30%. $\hat{R}$ refers to an improved version of the Gelman-Rubin convergence diagnostics. ESS is the effective sample size.

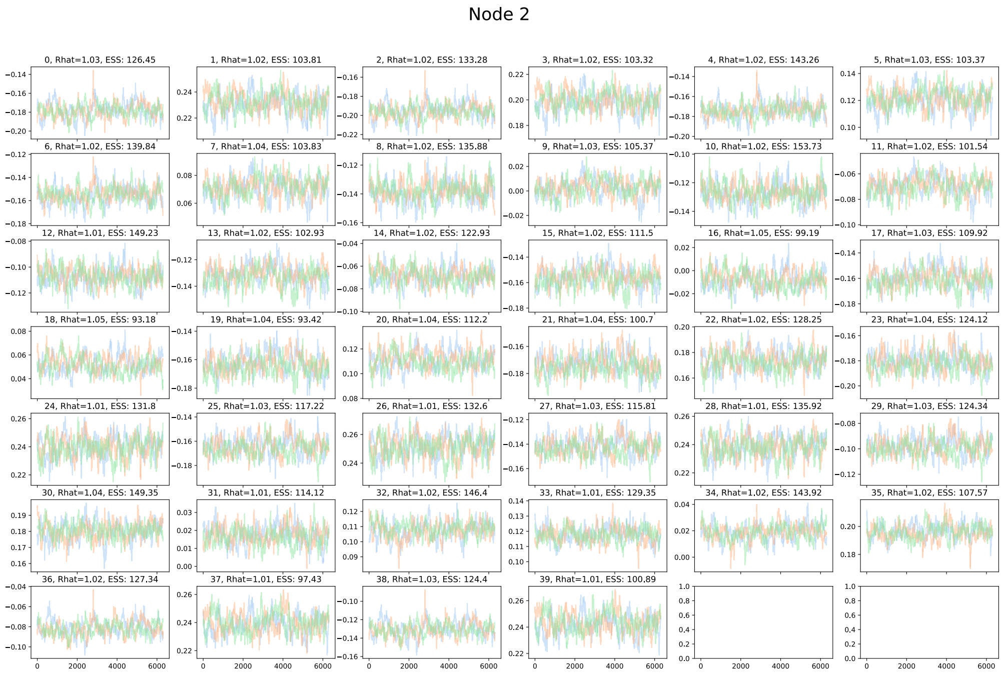


Figure 29: Trace plots for estimation of ancestral states. Colors indicate the result of three independent chains. Trace plots are shown after removing a burnin of 30%. $\hat{R}$ refers to an improved version of the Gelman-Rubin convergence diagnostics. ESS is the effective sample size.

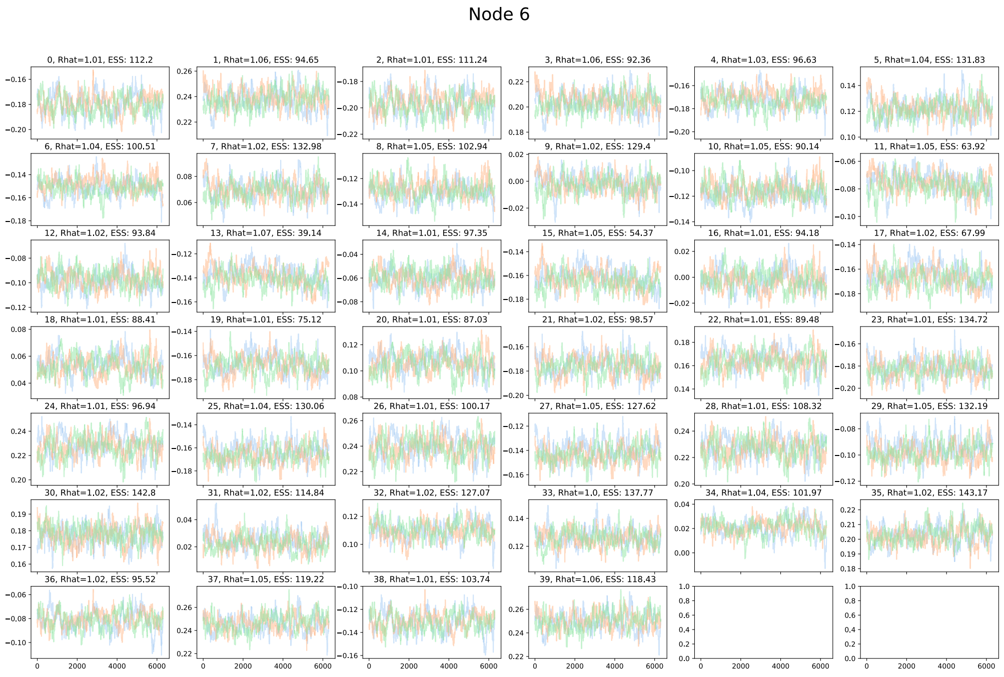


Figure 30: Trace plots for estimation of ancestral states. Colors indicate the result of three independent chains. Trace plots are shown after removing a burnin of 30%. $\hat{R}$ refers to an improved version of the Gelman-Rubin convergence diagnostics. ESS is the effective sample size.

### 7 Supplementary Material for section *"Future Directions"*

#### 7.1 Simulation

We simulate 20 datasets on the phylogeny given in Figure 4b in the main manuscript using the stochastic shape model with parameters $\alpha = 0.005$, $\delta = 0.05$ and $\sigma = 0.3$. The root shape used for simulation is a simulated butterfly forewing with centroid size 1 (see Fig. 31).

#### 7.2 Inference

We infer the posterior using the our metropolis-hastings algorithm with the following settings: $N = 3000$, $\delta = 0.05$, $\lambda = 0.95$, $\gamma = 0.0001$, $l_s = 1$, $\tau_\sigma = 0.1$ and $\tau_\alpha = 0.005$. We assume the following priors over the paramter $\sigma \sim \mathcal{U}(0, 1)$ and $\alpha \sim \mathcal{U}(0, 0.03)$. We use a mirrored Gaussian as proposal distribution and let the super root be the phylogenetic mean of our data.

#### 7.3 Convergence

We acheieve $\hat{R}$ values smaller than 1.1 for all dimensions of all landmarks as well as the parameters for all runs.

#### 7.4 Results

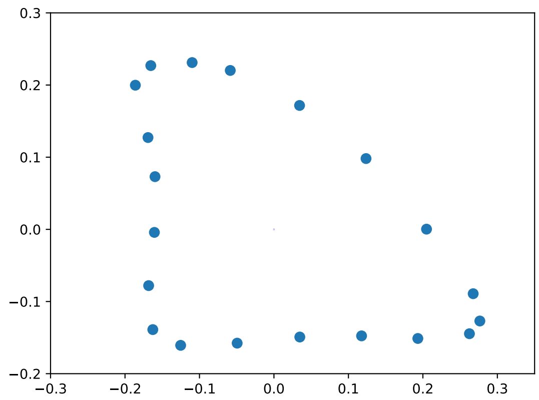


Figure 31: Root shape used for simulating data.

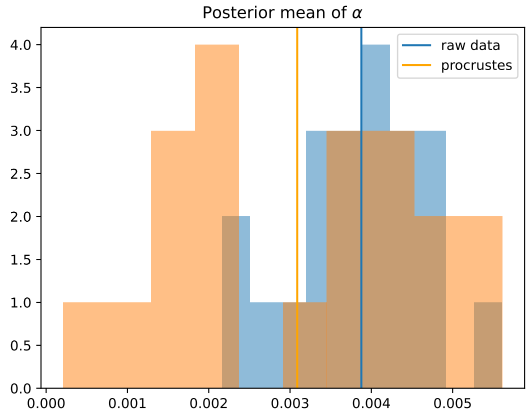


Figure 32: Posterior mean estimates for the $\alpha$ parameter.

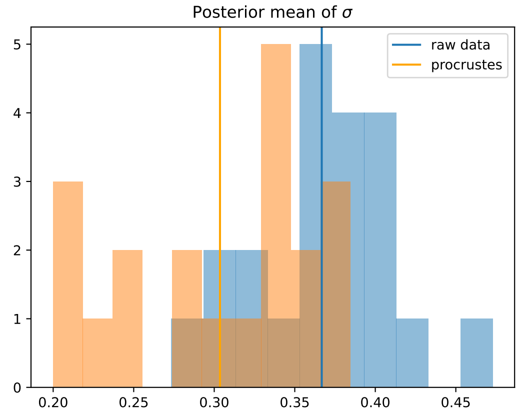


Figure 33: Posterior mean estimates for the $\sigma$ parameter.

### 8 Supplementary Material (extra) for section *"Root Estimation"*

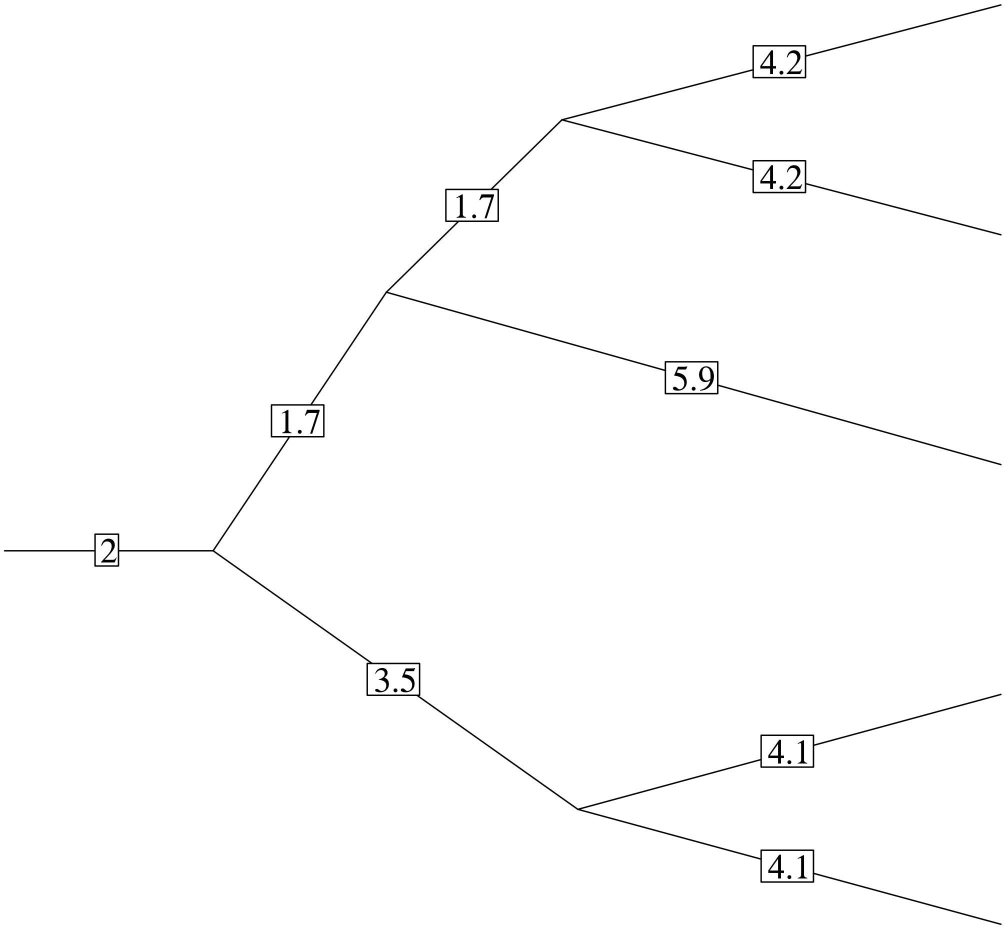


Figure 34: Mixed phylogeny used for inference.

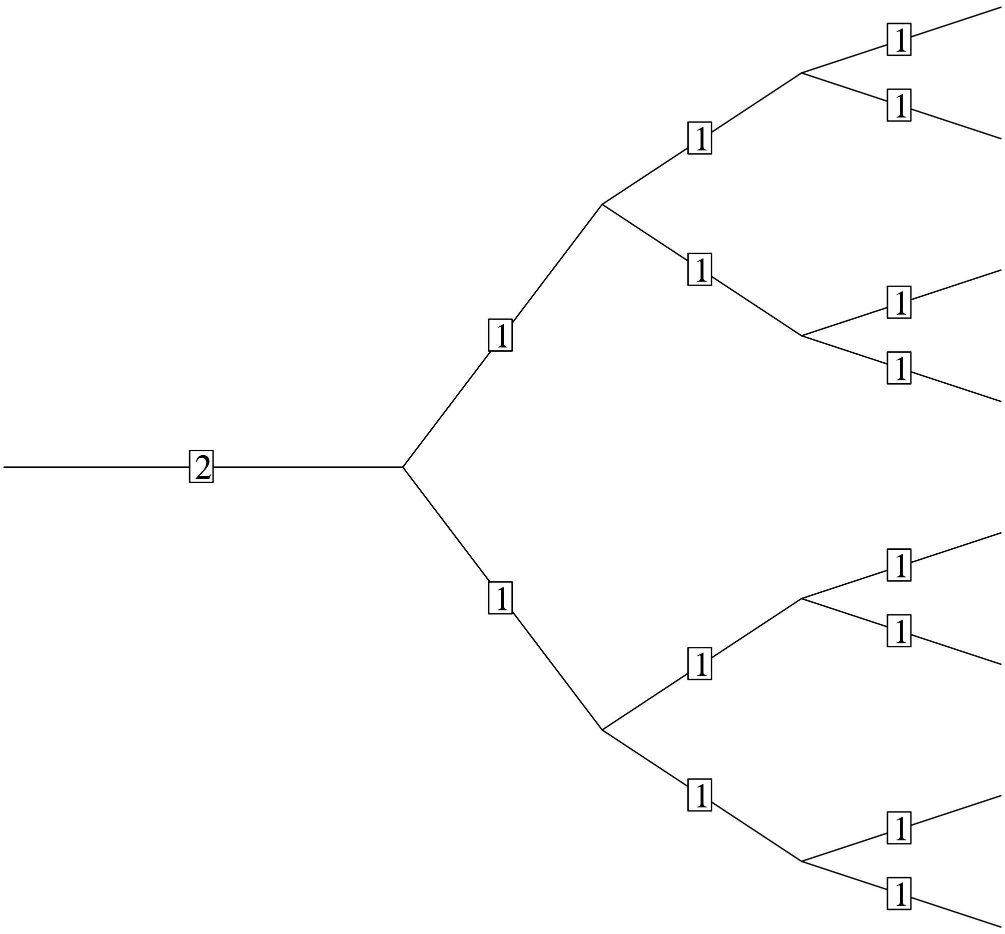


Figure 35: Symmetric phylogeny used for inference.

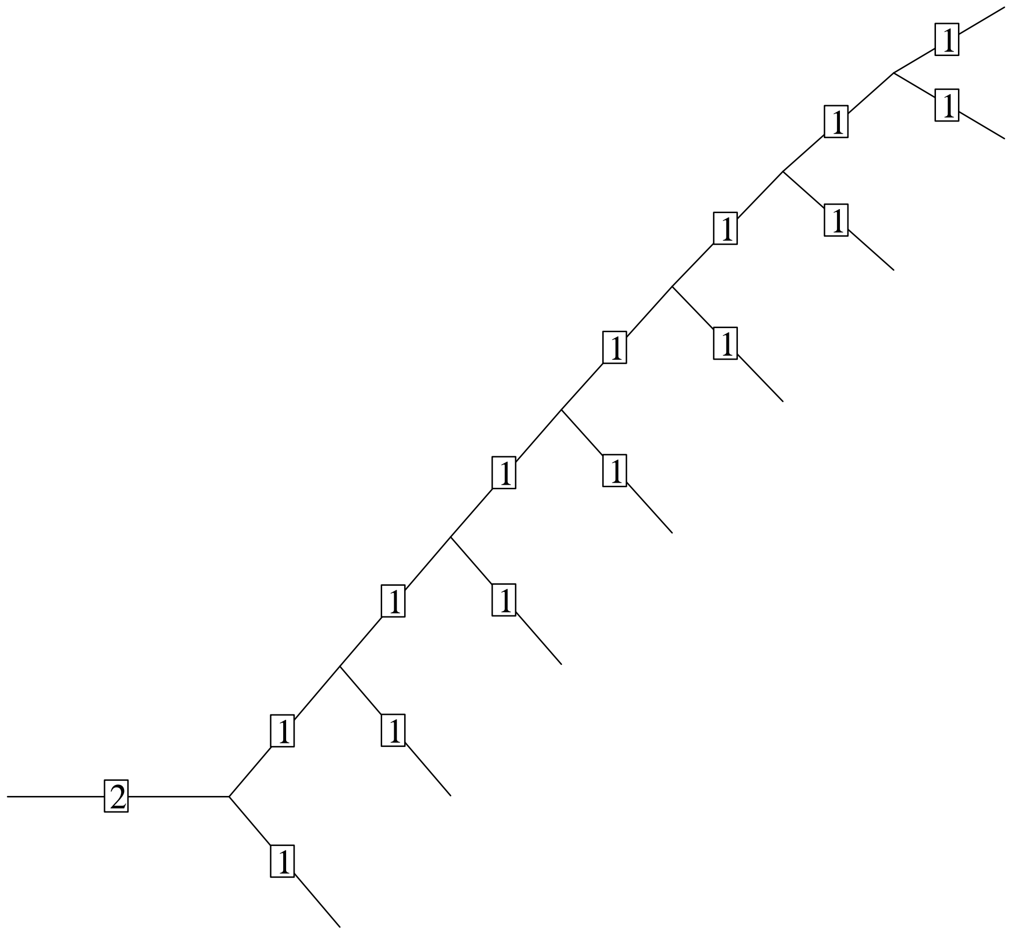


Figure 36: Asymmetric phylogeny used for inference.
